# CRISPR-mediated correction of skeletal muscle Ca^2+^ handling in a novel DMD patient-derived pluripotent stem cell model

**DOI:** 10.1101/2022.02.17.480850

**Authors:** Cristina Morera, Jihee Kim, Amaia Paredes-Redondo, Muriel Nobles, Denis Rybin, Robert Moccia, Anna Kowala, Jinhong Meng, Seth Garren, Pentao Liu, Jennifer E Morgan, Francesco Muntoni, Nicolas Christoforou, Jane Owens, Andrew Tinker, Yung-Yao Lin

## Abstract

Mutations in the *dystrophin* gene cause the most common and currently incurable Duchenne muscular dystrophy (DMD) characterized by progressive muscle wasting. Although abnormal Ca^2+^ handling is a pathological feature of DMD, mechanisms underlying defective Ca^2+^ homeostasis remain unclear. Here we generate a novel DMD patient-derived pluripotent stem cell (PSC) model of skeletal muscle with an isogenic control using clustered regularly interspaced short palindromic repeat (CRISPR)- mediated precise gene correction. Transcriptome analysis identifies dysregulated gene sets in the absence of dystrophin, including genes involved in Ca^2+^ handling, excitation-contraction coupling and muscle contraction. Specifically, analysis of intracellular Ca^2+^ transients and mathematical modeling of Ca^2+^ dynamics reveal significantly reduced cytosolic Ca^2+^ clearance rates in DMD-PSC derived myotubes. Pharmacological assays demonstrate Ca^2+^ flux in myotubes is determined by both intracellular and extracellular sources. DMD-PSC derived myotubes display significantly reduced velocity of contractility. Compared with a non-isogenic wild type PSC line, these pathophysiological defects could be rescued by CRISPR-mediated precise gene correction. Our study provides new insights into abnormal Ca^2+^ homeostasis in DMD and suggests that Ca^2+^ signaling pathways amenable to pharmacological modulation are potential therapeutic targets. Importantly, we have established a human physiology-relevant *in vitro* model enabling rapid pre-clinical testing of potential therapies for DMD.

## 1. Introduction

Skeletal muscle is responsible for voluntary movements and breathing. The dystrophin-glycoprotein complex (DGC) is enriched on the sarcolemma of muscle fibers and maintains muscle integrity by connecting the intracellular actin cytoskeleton with the extracellular matrix (ECM) ligands through the glycosylated cell-surface receptor dystroglycan [1]. In addition to its structural role, dystrophin recruits cytosolic components syntrophin, dystrobrevin, and neuronal nitric oxide synthase (nNOS) to the DGC, which is also involved in signaling pathways that regulate muscle homeostasis [2, 3]. Loss-of-function mutations in the dystrophin (*DMD*) gene cause the most common X-linked neuromuscular disorder, Duchenne muscular dystrophy (DMD), affecting about 1 in 3,500 to 5,000 boys. DMD is characterized by progressive weakness and wasting of skeletal muscle, followed by accumulation of fat and connective tissue. Current standards of care and emerging therapies for DMD can only delay the disease progression, but do not provide effective treatments. Consequently, DMD patients invariably lose ambulation and develop cardiorespiratory complications that severely limits life expectancy [4, 5].

The *DMD* gene is composed of 79 exons, spanning approximately 2.2 Mb on the X chromosome, and hence the largest human gene. To date, more than 7,000 mutations in the *DMD* gene have been reported to cause DMD [6]. The expression of *DMD* transcripts is under the control of 7 tissue-specific promoters that use unique first exons and splicing into distal exons, resulting in several dystrophin isoforms that differ in their N-terminus and length, including Dp427, Dp260, Dp140, Dp116 and Dp71 [5]. The full-length dystrophin isoform Dp427, encoded by a ∼14 kb cDNA expressed in muscle and brain, can be largely categorized into four domains: the N-terminal actin binding domain (ABD), the central rod domain (including a second ABD), the cysteine-rich domain and the carboxyl-terminal domain (Fig. 1A). Out-of-frame deletions and duplications are the most common mutations responsible for ∼75% of patients with DMD. In-frame deletions of the central rod domain cause Becker muscular dystrophy (BMD), which is the allelic and milder form of DMD [7]. It has been suggested that restoration of a shortened but partially functional dystrophin is a valid approach to ameliorate dystrophic muscle phenotypes [8]. Current therapeutic strategies for treating DMD include: clinically approved antisense oligonucleotide-mediated exon-skipping [9] and premature termination codon readthrough [10]; those in clinical trials, e.g. adeno-associated virus (AAV)-mediated micro-dystrophin gene therapy [11]; and those in pre-clinical development, e.g. CRISPR-mediated in-frame deletion of *DMD* exons [12]. However, it should be noted that restored expression of internally-deleted forms of dystrophin does not always guarantee their function because some in-frame deletions cause severe phenotypes in DMD patients while others lead to BMD phenotypes that are still associated with progression of weakness [5, 8]. To ensure clinical efficacy of the aforementioned strategies for treating DMD, the functional impact of different forms of shortened dystrophin requires extensive pre-clinical assessments.

**Fig. 1.**
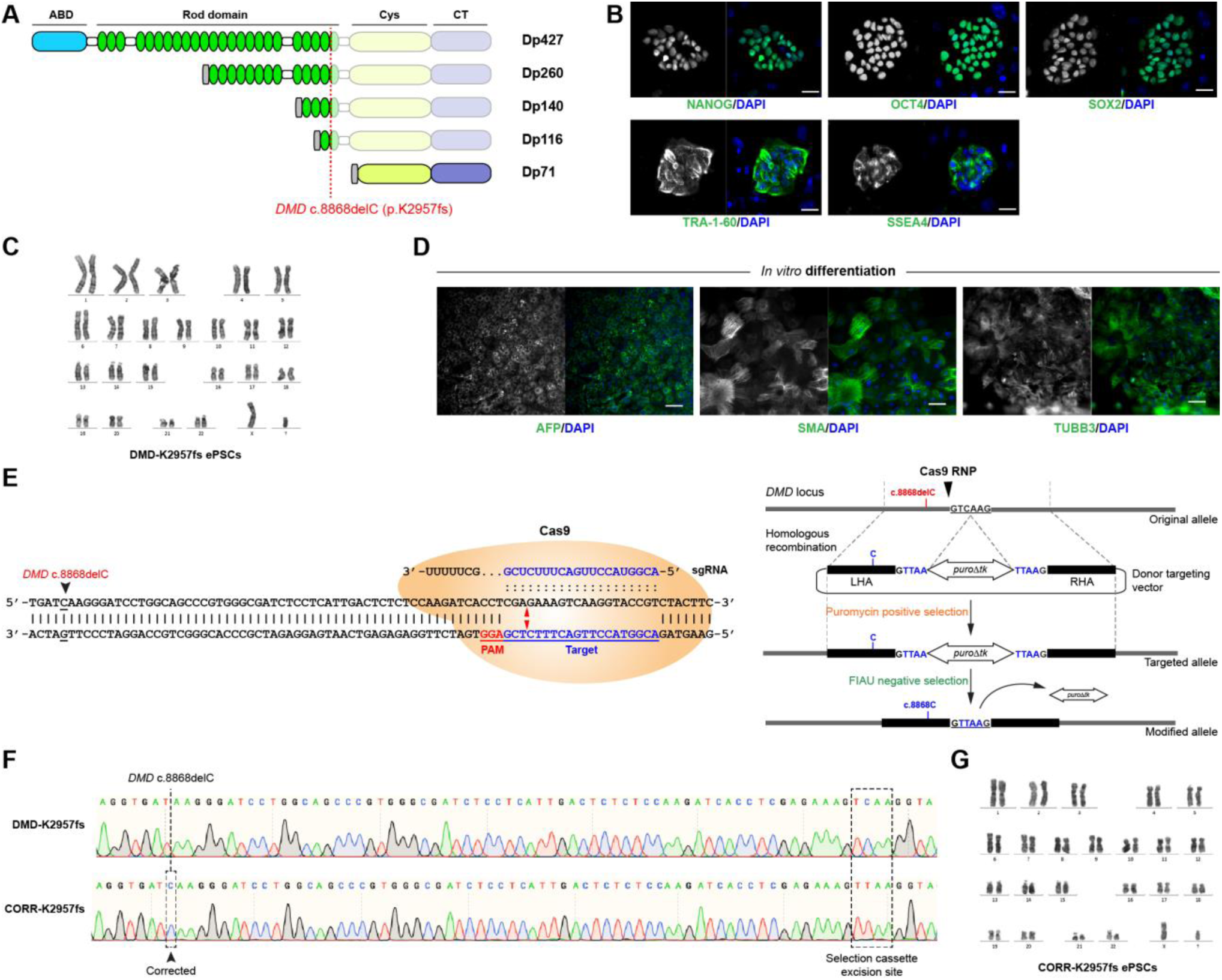
Generation of a novel DMD patient-derived PSC line with CRISPR-corrected isogenic control. (A) A schematic diagram shows dystrophin isoforms, including Dp427, Dp260, Dp140, Dp116 and Dp71. The full-length dystrophin isoform Dp427 contains four major domains: the N-terminal actin binding domain (ABD), the central rod domain (containing a second ABD), the cysteine-rich domain and the carboxyl-terminal domain. The *DMD* c.8868delC mutation affects all dystrophin isoforms, except Dp71. (B) Representative images of immunocytochemistry demonstrate that DMD-K2957fs ePSCs express specific pluripotency-associated markers, such as NANOG, OCT4, SOX2, Tra-1-60 and SSEA4. Scale bar, 30 μm. (C) DMD-K2957fs ePSCs have a normal karyotype. (D) *In vitro* differentiation of DMD-K2957fs ePSCs to cells representing three germ layers as demonstrated by immunocytochemistry of α-fetoprotein (AFP, endoderm), smooth muscle actin (SMA, mesoderm), and tubulin beta 3 class III (TUBB3, ectoderm). Scale bar, 100 μm. (E) A schematic diagram of the genome editing strategy that utilizes sgRNA and Cas9 RNP complex targeting specific site near the *DMD* c.8868delC mutation, stimulating homology-directed repair, followed by puromycin positive selection and FIAU negative selection to identify precisely corrected clones (CORR-K2957fs ePSCs). (F) Sanger sequencing demonstrates the precise correction of the *DMD* c.8868delC and the TTAA sequences at the selection cassette excision site in CORR-K2957fs ePSCs. (G) The karyotype of CORR-K2957fs ePSCs remains normal after CRISPR-mediated genome editing.

To elucidate pathological mechanisms underlying dystrophin deficiency and to evaluate potential therapeutic strategies, numerous animal models have been developed for DMD, including mice and larger animals, such as dogs and pigs [13, 14]. However, animal models may not be ideal for investigating the human pathophysiology of DMD and pre-clinical testing of candidate therapeutics. For example, the most commonly used dystrophin-deficient *mdx* mice have a near normal life span and do not show comparable disease severity in relation to human DMD patients [13, 14]. In addition, many therapeutic interventions that ameliorated phenotypes in the *mdx* mice have shown poor clinical translation to humans, e.g. the PDE5 inhibitors [15] and a utrophin modulator, ezutromid [16]. Although DMD patient primary myoblasts have been used to study the pathogenesis of DMD, the lack of isogenic controls and finite passage numbers have limited their applications in disease modeling and large-scale drug discovery. Thus, there is an unmet need for human physiologically-relevant DMD *in vitro* models of skeletal muscle with corresponding isogenic controls.

In terms of skeletal muscle physiology, the contraction of skeletal muscle is induced by the excitation-contraction (E-C) coupling process, reviewed in [17]. When the neurotransmitter acetylcholine binds to nicotinic acetylcholine receptors (ligand-gated ion channels) on the sarcolemma of muscle fibers, this leads to depolarization, voltage-gated sodium channel opening and spread of depolarization to the T-tubules (invagination of the sarcolemma). This causes a conformational change of the L-type voltage-dependent Ca^2+^ channel (Ca_V_1.1), which in turn activates the opening of ryanodine receptor (RYR) on the sarcoplasmic reticulum (SR) to release Ca^2+^ from the SR lumen into the cytoplasm. This leads to activation of actin-myosin sliding within the sarcomere (the contractile unit of a muscle fiber), resulting in muscle contraction. During the relaxation of skeletal muscle, sarcoplasmic/endoplasmic reticulum Ca^2+^-ATPase (SERCA) on the SR transfers cytosolic Ca^2+^ back to the lumen of SR. It has been long suggested that defective Ca^2+^ handling underlies the pathophysiology of DMD, leading to increased susceptibility to myofiber necrosis [3, 17]. Nevertheless, the molecular and cellular mechanisms underlying dysregulation of Ca^2+^ homeostasis in the absence of dystrophin remain poorly understood.

Recent studies have highlighted the opportunities and challenges of human pluripotent stem cell (PSC)-derived skeletal muscle as a disease model to develop novel therapies for muscular dystrophies [18, 19]. Here we have generated a novel DMD patient-derived PSC line carrying the *DMD* c.8868delC mutation and used CRISPR-mediated genome editing to precisely correct the *DMD* mutation generating a corresponding isogenic control line. Transcriptome analysis of the isogenic pair of PSC-derived myogenic cultures identified down-regulated gene sets in DMD, including genes involved in Ca^2+^ handling and E-C coupling such as *ATP2A1*, which encodes SERCA1. Comparative analysis of intracellular Ca^2+^ transients and mathematical modeling revealed that DMD-PSC derived myotubes have significantly reduced Ca^2+^ clearance rates, compared to wildtype or CRISPR-corrected isogenic controls. Consistent with transcriptome analysis, myotube contractility was also compromised in the absence of dystrophin. Together, our findings provide novel insights into mechanisms underlying defective Ca^2+^ homeostasis in DMD pathogenesis and demonstrate a human-relevant *in vitro* platform with functional readouts, which enables rapid pre-clinical assessment of potential therapies for treating DMD.

## 2. Materials and Methods

### 2.1 Ethical approval

Patient fibroblast lines were obtained from the MRC Centre for Neuromuscular Diseases Biobank (REC reference 06/Q0406/33). For the use of these cells, we have appropriate ethics approval (REC reference 13/LO/1826; IRAS project ID: 141100). In addition, all patients or their legal guardians gave written informed consent for industrial collaboration. The wildtype human BIONi010-C iPSC line was acquired from EBiSC (https://cells.ebisc.org).

### 2.2 Maintenance and myogenic differentiation of PSC lines

*DMD-K2957fs and CORR-K2957fs ePSCs*. Using a recently developed expanded potential stem cell medium (EPSCM) and a six-factor based reprogramming protocol [20, 21], we generated DMD- K2957fs patient-derived PSCs that could be stably maintained in the EPSCM, referred to as DMD- K2957fs ePSCs, as well as the CRISPR-corrected isogenic control CORR-K2957fs ePSCs. Mouse embryonic fibroblasts (MEFs) were used as feeder cells for co-culturing with the human ePSCs. MEFs were cultured in M10 medium, containing DMEM (Gibco, 10829), 10% Fetal Bovine Serum (Gibco, 10270-098), 1% Penicillin-Streptomycin-Glutamine (Gibco, 10378-016) and 1% Non-Essential Amino Acids solution (Gibco, 11140-035), and expanded before treating them 2.5 h with Mitomycin C (Sigma, M4287). After treated they were plated in M10 media in gelatin (Sigma, G-1393) coated plates (5 mins, 37°C) at a density of 7x10^4^ cells/cm^2^ 24 h before thawing the ePSCs. Human ePSCs were thawed at 37°C and plated on top of the feeder cells in EPSCM with 10 µM Y-27632 dihydrochloride (Tocris, 1254) for 48 h. When the cells were confluent, they were washed in 1X PBS and dissociated using Accutase (Millipore, SRC005) for 5 mins at 37°C. Accutase was neutralized with EPSCM with 10 µM Y-27632 and the cells were filtered with 100 µm strainers (Sysmex, 04-004- 2328). After centrifugation at 1,200 rpm for 5 mins, ePSCs were resuspended and plated at 3x10^4^ cells/cm^2^ in EPSCM with 10 µM Y-27632 for expansion.

*BIONi010-C iPSCs*. Vitronectin coated plates (Life Technologies, A14700, 1:100 in DPBS 1h at RT) were used to culture BIONi010-C iPSCs with StemFlex medium (Fisher scientific, 15627578) supplemented with Revitacell (Fisher Scientific, 15317447) for 24h. When the cells were confluent, they were washed in 1x DPBS and dissociated using Versane (Life technologies, 15040066) for normal clump passage and expansion.

*Transgene-free myogenic differentiation*. As described [22], human PSC lines were differentiated to myogenic progenitor cells (MPCs) within 3∼4 weeks and subcultured in skeletal muscle growth medium (Promocell, C-23260). Cryopreserved MPCs were thawed in skeletal muscle growth medium (GM) and induced to form multinucleated myotubes in N2 differentiation medium (DM). All medium recipes have been described [22].

### 2.3 CRISPR-mediated precise genome editing

The genome editing strategy used in this study was as described [23]. A donor targeting vector was constructed using Gibson Assembly (NEB, E2611S) to incorporate the *piggyBac* (*PGK-puroΔtk*) selection cassette flanked by two 1-kb homology arms specific to the *DMD* locus. A sgRNA containing the 5’-ACGGUACCUUGACUUUCUCG-3’ target sequence was synthesized by Synthego. The sgRNA was mixed with EnGen Cas9 NLS (NEB, M0646T) to form a ribonucleoprotein complex, then mixed with the donor targeting vector and DMD-K2957fs ePSCs, followed by electroporation using Lonza 4D.

### 2.4 Karyotype analysis

Prior to metaphase harvesting, ePSCs were grown in M15 medium (Knockout DMEM, 15% Fetal Bovine Serum, 1X glutamine-penicillin-streptomycin, 1X nonessential amino acids, 50 mM β- Mercaptoethanol and 1 ng/ml human recombinant leukemia inhibitory factor) for 24 h and then treated with 0.1 μg/ml Colcemid (Gibco, 15210-040) for 2.5 h. After harvesting, cells were treated with a hypotonic solution (0.075 M KCl) for 15min at 37°C and fixed with methanol:acetic acid (3:1) solution. Chromosomes were stained and analysed by G-banding. The karyotype analysis was performed by the Cytogenetics Service at the Barts Health NHS Trust.

### 2.5 Immunocytochemistry

Cultures were fixed with 4% paraformaldehyde (PFA) (SantaCruz, sc-281692) for 20 mins at RT. Before and after fixation, the cells were washed with 1X PBS 3 times. Samples were permeabilized with 0.5% Triton 100X (Sigma, T8787) in 1X PBS for 15 mins at RT. After 3 washes with 1X PBS, samples were blocked with 10% goat serum (Sigma, G9023) in 1X PBS for 1 hour at RT, followed by incubation with primary antibodies in blocking buffer at 4°C overnight. Dilution of primary antibodies are as following: NANOG (Abcam, AB80892; 1:100), OCT4 (Santa Cruz, sc-5279; 1:100), SOX2 (R&D, MAB2018; 1:100), Tra-1-60 (Santa Cruz, sc-21705; 1:100), SSEA4 (BD Bioscience, 560796; 1:50), α-fetoprotein (R&D, MAB1368; 1:100), α-smooth muscle actin (R&D, MAB1420; 1:75), tubulin beta 3 class III (R&D, MAB1195; 1:100), PAX7 (DSHB, 1:100), MYOD1 (Dako, M3512; 1:100), MYOG (DSHB, F5D; 1:100), MF20 (DSHB, 1:100), titin (DSHB, 9D10; 1:100), dystrophin (Millipore, MABT827; 1:50). The next day samples were washed with 1X PBS for 3 times, followed by incubation with appropriate Alexa Fluor secondary antibodies (1:500) and DAPI (1:500) (Sigma, D9542) in blocking buffer for 1 hour at RT. After wash, the samples were kept in 1X PBS at 4°C in the dark until analysis.

### 2.6 Immunoblotting

Protein lysates were collected using Radio-Immunoprecipitation Assay (RIPA) buffer (Sigma), supplemented with protease inhibitor (Roche) on ice for 15 min. The lysate was then boiled for 5 min and centrifuged at 14,000 x g for 10 minutes at 4°C. Protein concentration was determined using a Pierce™ BCA Protein Assay Kit (Thermo Fisher). 30µg/well of each sample was loaded onto NuPAGE Novex 3-8% Tris-Acetate gel (Thermo Fisher), and proteins were separated at a constant voltage of 150V for 1.5 hours, before being transferred to Polyvinylidene difluoride (PVDF) membrane using a Trans-Blot Turbo Transfer system (Bio-rad). The membrane was blocked with Odyssey block solution (LI-COR Biosciences, Cambridge, UK) for one hour, and then incubated with primary antibodies against dystrophin (rabbit polyclonal IgG (H+L), 1:2000; Fisher Scientific, PA5- 32388), using GAPDH (mouse monoclonal IgG1, 1:5,000; Thermo Fisher) as housekeeping control. After washing with PBS containing 0.1% Tween 20 (PBST) for 15 min, 3 times at RT, the membrane was incubated with IRDye 680RD goat anti-rabbit and IRDye 800CW goat anti-mouse secondary antibodies (1:15000, LI-COR Biosciences, Cambridge, UK) for 1 hour at RT. The image of the blotted membrane was acquired by Odyssey Clx infrared imaging system (LI-COR Biosciences, Cambridge, UK) using Image Studio Lite 5.2 software.

### 2.7 Calcium imaging

PSC-derived MPCs were seeded in glass bottom dishes (MatTek, P35G-1.5-14-C) at 80x10^3^ cells/cm^2^ during 5 days in N2 media. 50 µl of Pluronic F-127 solution (Molecular Probes, 10767854) was added to 50 µg of Fluo-4 AM (Molecular Probes, 11504786). Myotubes were loaded for 1h at 37°C with 10 µl of Pluronic/Fluo-4 in 1 ml of recording buffer. Recording buffer contained 150 mM NaCl, 10 mM HEPES, 2.6 mM KCl, 10 mM D-(+) glucose, 2mM CaCl_2_, 1 mM MgCl_2_, pH 7.4. Cells were rinsed with recording buffer before imaging with the 488 nm laser line of a confocal microscope and an emission filter of LD Plan-Neufluar 40x/0.6 korr (Zeiss LSM 880). To stimulate intracellular Ca^2+^ flux, 100 ul of 3M acetylcholine chloride (Sigma, A2661) was dissolved in recording buffer and added into the MatTek dish containing 1 ml buffer (final concentration 272.73 mM). Ca^2+^ release was recorded as a time series. Ca^2+^ release inhibition experiments were performed with 10 µM nifedipine (Sigma, N7634), 10 µM ryanodine (Santa Cruz, sc-201523), 10 µM cyclopiazonic acid (Sigma, C1530). Images were analyzed by ImageJ, drawing ROIs inside the myotubes and extracting its mean values of fluorescence, indicative of the Ca^2+^ transient. Using these data, we analyzed the F1/F0 (fluorescence peak/basal fluorescence).

### 2.8 Particle image velocimetry (PIV) analysis

Videos acquired from myogenic cultures stimulated by acetylcholine were analyzed using Particle Image Velocimetry in MATLAB software [24]. Image data were calibrated using a defined scale.

### 2.9 RNA sequencing

Total RNA was quality controlled on a Tapestation 4200 with a RIN cutoff of 7.0. RNAseq libraries were generated using the Illumina Truseq Stranded mRNA LP kit (20020594) and sequenced on a Nextseq 500 at 2x75 paired end using the NSQ 500/550 Hi Output KT v2.5 (150 CYS) (20024907). Libraries were generated on a Perkin Elmer Janus G3 automated system in batches. Each batch contained half of the replicates of each condition to control for batch effects. All samples were multiplexed and split into two sequencing runs. The RNA sequencing data has been deposited to Gene Expression Omnibus (accession number GSE189053).

### 2.10 Transcriptome analysis

Quality of fastq files was assessed using FastQC (v0.11.8) (http://www.bioinformatics.babraham.ac.uk/projects/fastqc/). Reads were aligned to the human reference genome (hg38, Ensembl version 98) using STAR (v2.7.3a) with additional settings: -- quantMode TranscriptomeSAM; --alignIntronMin 20; --alignIntronMax 1000000; -- alignSJoverhangMin 8; --alignSJDBoverhangMin 1; --alignMatesGapMax 1000000; --sjdbScore 1; --outFilterMultimapNmax 20; --outFilterMismatchNmax 1000; --outFilterMismatchNoverLmax 0.10 [25]. Additional quality control was assessed using PicardTools CollectRNASeqMetrics (http://broadinstitute.github.io/picard). Estimated counts were obtained using Salmon (v0.14.2) alignment-based mode with the --gcBias flag [26]. Differential expression analysis was performed using DESeq2 (v1.24.0) in R (v3.6.2) [27]. Genes with very low mean read counts across all samples were removed prior to fitting. The statistical model included a term for treatment group and processing batch. Shrinkage of fold-change estimates was performed using ashr (v2.2-32) [28]. Multiple comparisons were adjusted using the Benjamini-Hochberg method [29]. Gene set enrichment analysis was performed using the GSEA pre-ranked algorithm implemented in the R package fgsea using MsigDB Hallmark collection v6.2 as the input gene sets [30–33]. Genes were ranked using the post-shrinkage log2-fold-change estimates from DESeq2 analysis. Any gene that was excluded by the DESeq2 independent filtering algorithm was also excluded from GSEA analysis.

### 2.11 Real-time quantitative PCR (qPCR)

Complementary DNA (cDNA) was generated using High-Capacity cDNA Reverse Transcription Kit (Applied Biosystems, 4368814), followed by real-time quantitative PCR using PowerUp SYBR Green Master Mix (Applied Biosystems, A25742). The reactions were set up as per the manufacturer’s instructions and the primers are specified in Table S6. Cycle Thresholds were produced using StepOne Plus software (Applied Biosystems), normalized to the values of the reference gene *ACTB* [34], and relative gene expression levels were plotted as fold changes.

## 3. Results

### 3.1 Generation of a novel DMD patient-derived pluripotent stem cell line

We used a dermal fibroblast line carrying the *DMD* c.8868delC (p.K2957fs) mutation in exon 59 from a male patient diagnosed with DMD. The clinical features of this patient were those typical of DMD with delayed motor milestones, bilateral calf hypertrophy, proximal muscle weakness, positive for Gower’s sign, and markedly elevated serum levels of creatine kinase (>15,000 IU/L). The patient’s muscle biopsy report confirmed dystrophic features, absence of the dystrophin protein, as well as reduced sarcoglycans (Table S1). The *DMD* c.8868delC mutation is predicted to cause a frameshift and premature termination of translation, affecting dystrophin isoforms Dp427, Dp260, Dp140, Dp116, but not Dp71 (Fig. 1A). Using a recently developed expanded potential stem cell medium (EPSCM) and a six-factor based reprogramming protocol [20, 21], we generated DMD-K2957fs patient-derived pluripotent stem cells (PSCs) that can be stably maintained in the EPSCM, referred to as DMD- K2957fs EPSCM-PSCs (hereinafter ePSCs). Gene expression analysis confirmed *OCT4* and *NANOG* mRNA expression, compared with parental fibroblasts (Fig. S1). Immunocytochemistry demonstrated that DMD-K2957fs ePSCs expressed pluripotency markers, including NANOG, OCT4, SOX2, Tra- 1-60 and SSEA-4 (Fig. 1B). The established DMD-K2957fs ePSC line had a normal karyotype (46, XY) (Fig. 1C). Finally, *in vitro* differentiation confirmed that DMD-K2957fs ePSCs were capable of forming embryoid bodies and differentiating into cell types representing the three embryonic germ layers, as demonstrated by lineage-specific markers, β-III tubulin (ectoderm), smooth muscle actin (mesoderm) and α-fetoprotein (endoderm) (Fig. 1D). These results suggest that we have generated a novel patient-derived PSC line for DMD.

### 3.2 Precise correction of the *DMD* mutation by CRISPR-mediated genome editing

To precisely correct the *DMD* c.8868delC mutation in the DMD-K2957fs ePSC line, we employed a genome editing strategy [23], utilizing homology directed repair (HDR) stimulated by CRISPR/Cas9 site-specific endonuclease. First, we identified and synthesized an appropriate single guide RNA (sgRNA) targeting 56 bp downstream of the *DMD* c.8868delC mutation (Fig. 1E; Table S2). By delivering the sgRNA-Cas9 ribonucleoproteins, DNA double-strand breaks were introduced near the *DMD* mutation (Fig. 1E). To stimulate HDR, a donor targeting vector was delivered simultaneously. The donor targeting vector consists of two 1-kb homology arms flanking a *piggyBac* (*PGK-puroΔtk*) selection cassette (Fig. 1E). Note that the *DMD* mutation was corrected on the left homology arm and the nucleotide sequences at the junctions between either homology arm and the selection cassette were modified to accommodate the selection cassette excision site, TTAA (Fig. 1E). By positive selection with puromycin in EPSCM, we identified targeted clones that had an integrated donor vector by rapid PCR genotyping (Fig. S1) and confirmed that by sequencing. Next, the *piggyBac* (*PGK-puroΔtk*) selection cassette was removed by negative selection. To do this, targeted clones were electroporated with a plasmid expressing *piggyBac* transposase, followed by negative selection in EPSCM containing 1-(2-Deoxy-2-fluorob- D-arabinofuranosyl)-5-iodo-2,4(1H,3H)-pyrimidinedione (FIAU). If cells retained the *piggyBac* (*PGK-puroΔtk*) selection cassette, FIAU would be processed to metabolites which were toxic to the cells. Finally, PCR genotyping identified CRISPR-corrected clones that had the selection cassette removed without re-integration (Fig. S1). Sequencing confirmed that the *DMD* c.8868delC mutation was precisely corrected (Fig. 1F) and the nucleotide sequences spanning the selection cassette excision site encode the same amino acids (Fig. 1F). Further characterization confirmed that the established CRISPR-corrected clonal line had a normal karyotype (46, XY) (Fig. 1G). Sequencing of top 5 predicted off-target sites of this line did not identify any undesired mutations (Table S2). Together, these results indicate that we have successfully generated a CRISPR-corrected isogenic control (hereinafter CORR-K2957fs) for the DMD-K2957fs ePSC line.

To address inter-individual variability, we acquired a healthy donor-derived human induced pluripotent stem cell (iPSC) line BIONi010-C from the European Bank for Induced pluripotent Stem Cells (EBiSC). Immunocytochemistry showed that the BIONi010-C iPSC line expressed classic pluripotency markers (Fig. S1). Thus, this human iPSC line could be used as a non-isogenic wildtype control for the DMD-K2957fs and CORR-K2957fs ePSC lines.

### 3.3 Transgene-free myogenic differentiation and restoration of full-length dystrophin expression

Using an efficient transgene-free myogenic differentiation protocol [22], we generated myogenic progenitor cells (MPCs) from the isogenic pair of CORR-K2957fs and DMD-K2957fs ePSC lines, as well as from the wildtype BIONi010-C iPSC line. This process was designated as primary myogenic differentiation. These PSC-derived MPCs were subcultured and cryopreserved (Fig. 2A). Upon culturing in skeletal muscle growth medium (GM) and then switching to differentiation medium (DM), the human PSC-derived MPCs elongated and fused to form multinucleated myotubes, while maintaining a pool of MPCs. This process was defined as secondary myogenic differentiation (Fig. 2A). Immunocytochemistry characterization of MPCs differentiated for 5 days demonstrated classic myogenic markers for all three lines, including MPCs expressing transcription factors PAX7, MYOD1, MYOG (Fig. 2B) and myotubes expressing sarcomere components myosin heavy chain (MF20) and titin (TTN) (Fig. 2B).

**Fig. 2.**
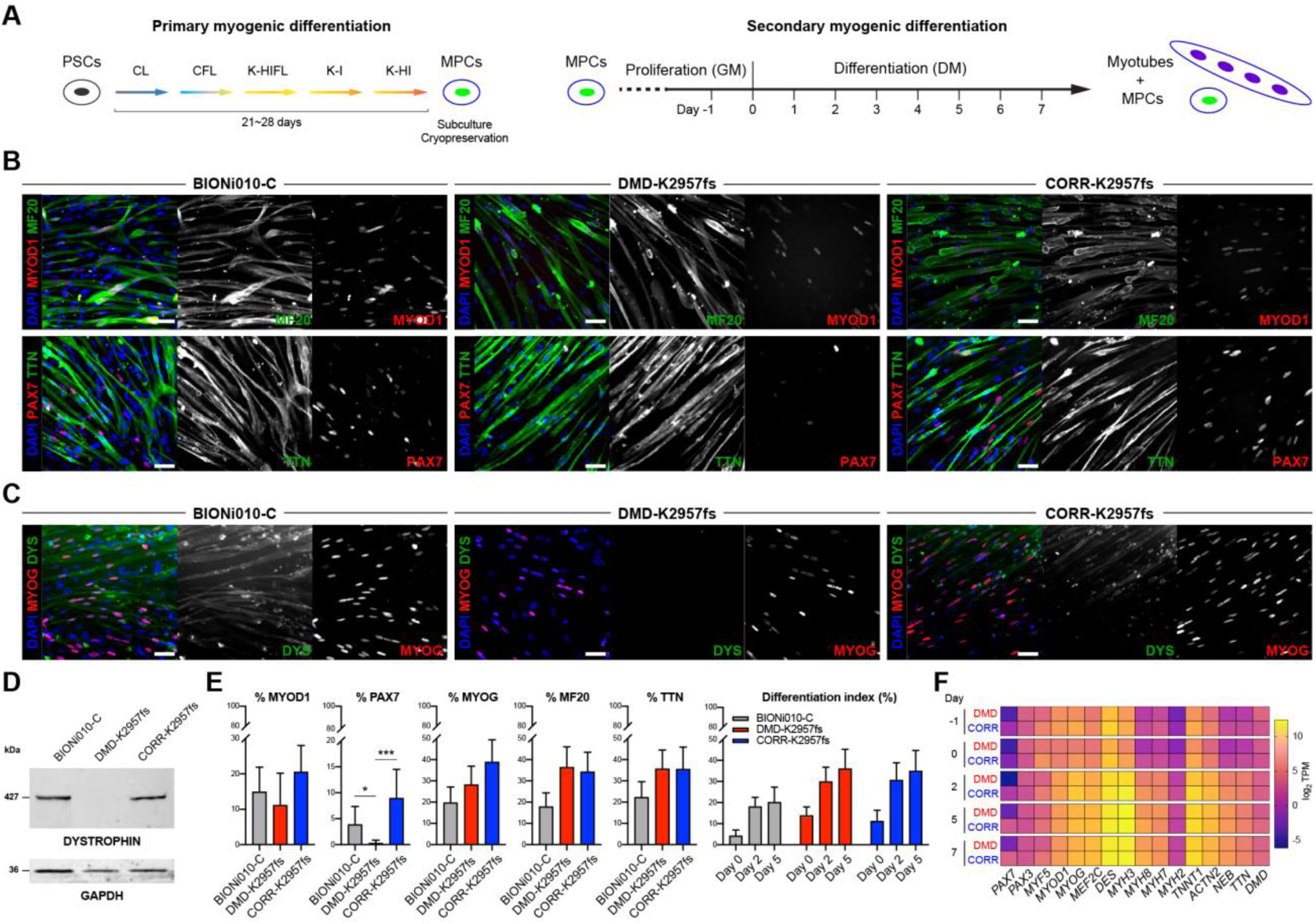
Characterization of transgene-free myogenic differentiation and restoration of dystrophin expression by CRISPR-mediated gene correction. (A) A schematic timeline of myogenic differentiation from human PSCs to MPCs (primary myogenic differentiation). Specific medium changes are indicated as CL, CFL, K-HIFL, K-I and K-HI. Human PSC-derived MPCs are expanded in growth medium (GM), which is switched to differentiation medium (DM) to induce fusion, resulting in multinucleated myotubes (secondary myogenic differentiation). (B) Representative images of Day 5 myogenic cultures derived from secondary differentiation of BIONi010-C, DMD-K2957fs and CORR-K2957fs MPCs. Immunocytochemistry demonstrates multinucleated myotubes in all three lines as shown by expression of myosin heavy chain (MF20) and sarcomeric protein titin (TTN), as well as myogenic transcription factors MYOD1 and PAX7. Scale bar, 50 μm. (C) Immunocytochemistry demonstrated that dystrophin protein expression (DYS) is completely absent in DMD-K2957fs myotubes (Day 5 cultures), and yet restored in CORR-K2957fs myotubes, similar to BIONi010-C myotubes. All three lines express myogenic transcription factor MYOG. Scale bar, 50 μm. (D) Immunoblotting confirmed that the full-length dystrophin (427 kDa) was present in BIONi010-C and CORR-K2957fs, but not detected in DMD-K2957fs myogenic cultures. GAPDH was used as the loading control. (E) Quantification of the percentage of MYOD1- and MYOG-positive MPCs, as well as MF20- and TTN-positive myotubes in BIONi010-C, DMD-K2957fs and CORR-K2957fs myogenic culture (Day 5). Note that the percentage of PAX7-positive cells is significantly lower in DMD-K2957fs myogenic culture compared to CORR-K2957fs and BIONi010-C. The differentiation index shows a similar trend between DMD-K2957fs and CORR-K2957fs myogenic cultures. Data are mean ± SD (n= 3; * p < 0.05, *** p < 0.001). (F) A heatmap of selected muscle-specific genes shows similar transcript expression patterns between DMD-K2957fs and CORR-K2957fs myogenic cultures across each time point of secondary myogenic differentiation, except *PAX7* transcript levels. TPM, Transcripts Per Million. Color scale represents log_2_ TPM.

Next, immunocytochemistry confirmed that the wildtype BIONi010-C iPSC-derived myotubes expressed dystrophin (Fig. 2C), whereas dystrophin expression was absent in DMD-K2957fs ePSC-derived myotubes (Fig. 2C). Importantly, CRISPR-mediated genome editing restored the dystrophin protein expression in the CORR-K2957fs ePSC-derived myotubes (Fig. 2C). Immunoblotting confirmed the presence of full-length dystrophin (427 kDa) in the BIONi010-C and CORR-K2957fs myogenic cultures, but not DMD-K2957fs (Fig. 2D).

On Day 5 of secondary differentiation, quantification of the percentage of MYOD1, MYOG, MF20 and TTN suggest that DMD-K2957fs and CORR-K2957fs ePSC derived myogenic cultures were very similar, while the percentage of PAX7-positive cells were significantly lower in DMD-K2957fs ePSC derived myogenic culture (Fig. 2E). The differentiation index, defined as the average of percentage of MF20 and TTN, also showed a similar trend between DMD-K2957fs and CORR-K2957fs myogenic cultures during secondary differentiation (Fig. 2E). Nevertheless, DMD-K2957fs displayed a compromised fusion competence (Fig. S2A), consistent with previous studies [35, 36]. We then performed RNA sequencing using cultures of DMD-K2957fs and CORR-K2957fs myogenic differentiation (Day -1, 0, 2, 5 and 7) and plotted the transcript levels of selected genes as a heatmap (Fig. 2F). DMD-K2957fs and CORR-K2957fs myogenic cultures expressed similar profiles of muscle-specific genes, such as transcription factors *MYF5*, *MYOD1*, *MYOG* and *MEF2C*, and sarcomeric components, including myosin heavy chain (MYH) isoforms *MYH3* (embryonic), *MYH8* (perinatal), *MYH2* (fetal), *DES, TNNT1*, *ACTN2*, *NEB* and *TTN*. Our transcriptome analysis did not detect expression of *MYH1* (late fetal) and *MYH4* (postnatal) transcripts (Supplementary Data S1), suggesting that human PSC-derived myotubes resemble fetal-like phenotypes. Note that *PAX7* was differentially expressed at each time point (Fig. 2F), suggesting that the transcript levels between DMD-K2957fs and CORR-K2957fs ePSC-derived myogenic cultures largely reflected the immunocytochemistry results.

### 3.4 Transcriptome analysis identifies affected biological processes in the absence of dystrophin

To further elucidate the differences between DMD-K2957fs and CORR-K2957fs myogenic cultures, we analyzed their transcriptomes. We first performed principal component analysis (PCA) to have an overview of intrinsic variability between samples. Plotting of the first two components (PC1, 63% variance and PC2, 9% variance) showed that sample replicates clustered together and segregated according to their differentiation time and genotypes, reflecting the direction of secondary myogenic differentiation from MPCs to multinucleated myotubes (Fig. 3A).

**Fig. 3.**
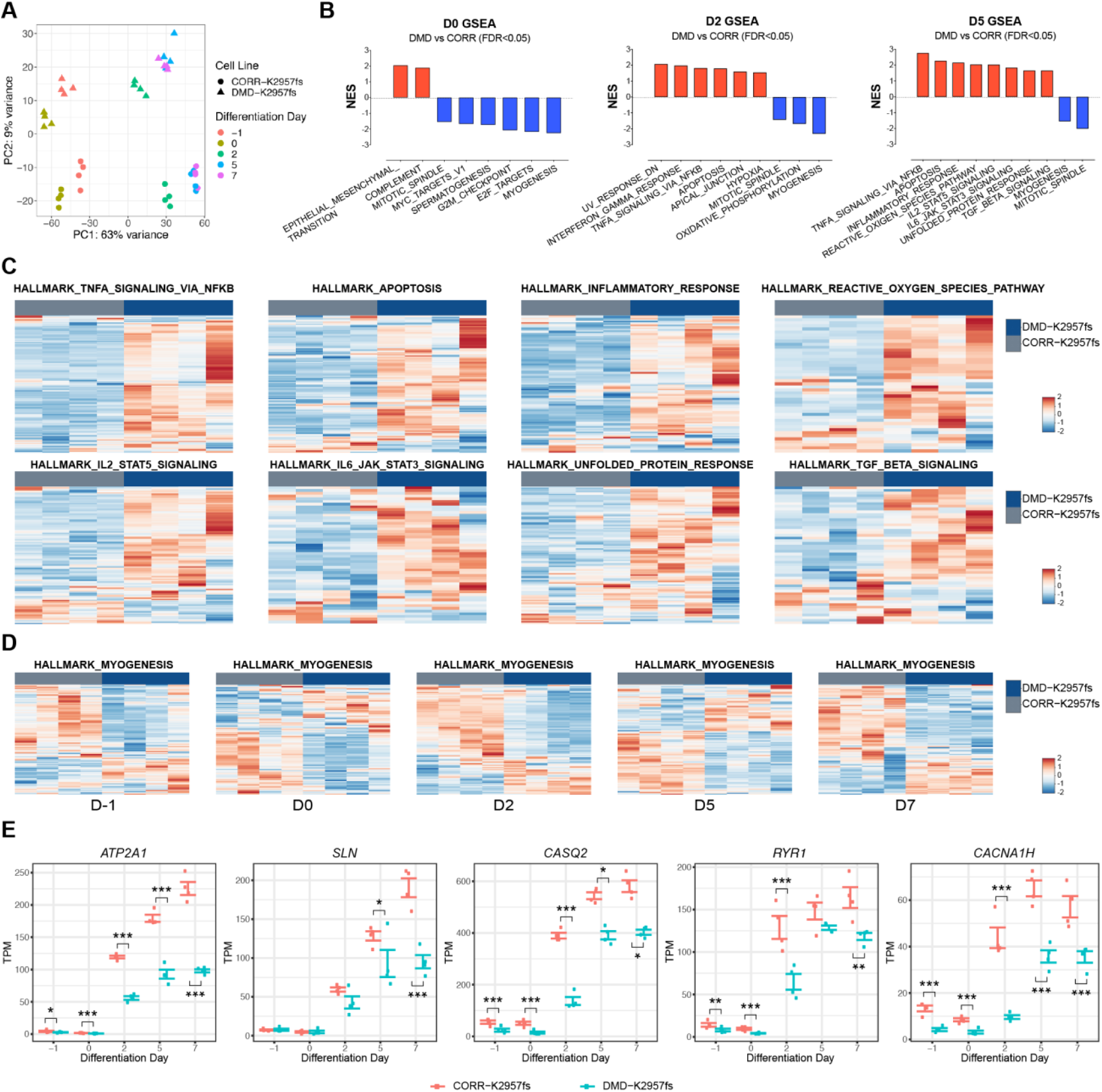
Transcriptome analysis reveals differentially regulated gene sets between DMD-K2957fs and CORR-K2957fs myogenic cultures. (A) A plot of principal component analysis of samples of two genotypes from secondary myogenic differentiation at Day -1, 0, 2, 5 and 7. Each condition has 4 biological replicates. (B) Selected HALLMARK gene sets that are significantly enriched in DMD-K2957fs (red bars) or CORR-K2957fs (blue bars) myogenic cultures at Day 0, 2 and 5 of secondary differentiation. NES, normalized enrichment score. FDR, false discovery rate. (C) Heatmaps of selected HALLMARK gene sets that are significantly enriched in DMD-K2957fs, compared to CORR-K2957fs myogenic cultures at Day 5. Color scales represent Z-scores. (D) Heatmaps of the HALLMARK_MYOGENESIS gene set that is significantly enriched in CORR- K2957fs myogenic culture, compared to DMD-K2957fs at all 5 time points. Color scales represent Z- scores. (E) Time-course expression levels of genes involved in Ca^2+^ handling within the HALLMARK MYOGENESIS gene set, including *ATP2A1*, *SLN*, *CASQ2*, *RYR1*, and *CACNA1H*. Wald test with Benjamini-Hochberg method for multiple comparison adjustment (* padj < 0.05, ** padj < 0.01, *** padj < 0.001). TPM, Transcripts Per Million. Each condition has 4 biological replicates.

Next, we examined the differential gene expression between the DMD-K2957fs and CORR-K2957fs transcriptomes at 5 timepoints. Differentially expressed (DE) genes between DMD-K2957fs and CORR-K2957fs were identified using DESeq2 [27, 28] (Supplementary Data S2) and visualized using volcano plots (Fig. S2B-F). To identify affected biological processes in the absence of dystrophin during myogenic differentiation, we performed Gene Set Enrichment Analysis (GSEA) [30, 31]. During secondary myogenic differentiation (DMD-K2957fs versus CORR-K2957fs comparison), the number of positively or negatively enriched HALLMARK gene sets with false discovery rate (FDR) <0.05 increased from ∼10 to more than 30 gene sets (Table S3; Supplementary Data S3). HALLMARK gene sets with positive normalized enrichment score (NES) indicated overall higher transcript levels in the DMD-K2957fs phenotype relative to CORR-K2957fs. Interestingly, many of these gene sets reflected several known DMD pathophysiological features (Table S3), such as TNFA SIGNALING VIA NFKB, APOPTOSIS, and INFLAMMATORY RESPONSE (Day 2, 5 and 7); IL6 JAK STAT3 SIGNALING, UNFOLDED PROTEIN RESPONSE, IL2 STAT5 SIGNALING, and REACTIVE OXYGEN SPECIES PATHWAY (Day 5 and 7); TGF BETA SIGNALING (Day 5) (Fig. 3B,C). In contrast, the MYOGENESIS gene set had significant negative NES at all time points, Day -1, 0, 2, 5 and 7 (Fig. 3D), indicating overall higher transcript levels in the CORR-K2957fs phenotype. This suggests that some genes within the MYOGENESIS gene set might have compromised function in DMD-K2957fs during myogenic differentiation. We investigated the “leading-edge” subset of the HALLMARK MYOGENESIS gene set that accounts for the enrichment signal and noticed that several Ca^2+^ handling related genes were significantly down-regulated in the DMD-K2957fs myogenic cultures, such as *ATP2A1*, *SLN, CASQ2*, *RYR1*, and *CACNA1H* (Fig. 3E). *SLN* encodes sarcolipin (a SERCA regulator); *CASQ2* encodes calsequestrin 2 (a high affinity Ca^2+^ binding protein in the terminal cisternae of SR); *CACNA1H* encodes Cav3.2 (a subunit in the voltage-gated T-type calcium channel). Consistent with transcriptome analysis (Fig. 3E), real-time qPCR showed that *ATP2A1*, *CASQ2*, *SLN*, *RYR1*, and *CACNA1H* were significantly down-regulated in DMD-K2957fs in comparison to CORR- K2957fs, and there was no significant difference for *ATP2A2* expression (Fig. S3A). While most of the above Ca^2+^ handling genes regulate E-C coupling, it is noteworthy that *CACNA1S* (Ca_V_1.1), not a member of the HALLMARK MYOGENESIS gene set but responsible for activating RYR opening to release Ca^2+^ from the SR, was significantly down-regulated in DMD-K2957fs compared to CORR- K2957fs at all 5 timepoints (Supplementary Data S2). Altogether, our findings suggest that Ca^2+^ homeostasis and E-C coupling in human PSC-derived myotubes might be affected in the absence of dystrophin.

### 3.5 Analysis of intracellular Ca^2+^ transients shows abnormal Ca^2+^ handling in DMD-K2957fs myotubes and CRISPR-mediated correction

We sought to investigate the regulation of Ca^2+^ homeostasis in skeletal muscle under healthy and DMD conditions. To do this, we performed calcium imaging in human PSC-derived myotubes to examine their Ca^2+^ handling properties. As previously described [37], the human PSC-derived myotubes were loaded with fluorescence-based Ca^2+^ indicator Fluo-4 and monitored using a laser scanning confocal microscope to examine the Ca^2+^ transients. Based on a previous publication [38], we determined the optimal acetylcholine concentration that gave stimulus-induced intracellular Ca^2+^ signals. Under acetylcholine stimulation, we observed consistent and robust cytosolic Ca^2+^ release and reuptake in human PSC-derived myotubes (Fig. 4A-C). The Ca^2+^ transient kinetics were then analyzed and compared between BIONi010-C, DMD-K2957fs, and CORR-K2957fs myotubes (Fig. 4A-C). Analysis of the basic properties of Ca^2+^ transients showed that BIONi010-C myotubes have significantly higher mean value of normalized Ca^2+^ transient amplitude (F1/F0) than DMD-K2957fs or CORR-K2957fs myotubes (Fig. 4D). Although the mean value of F1/F0 in CORR-K2957fs myotubes was higher than that in DMD-K2957fs myotubes, the difference was not statistically significant (Fig. 4D).

**Fig. 4.**
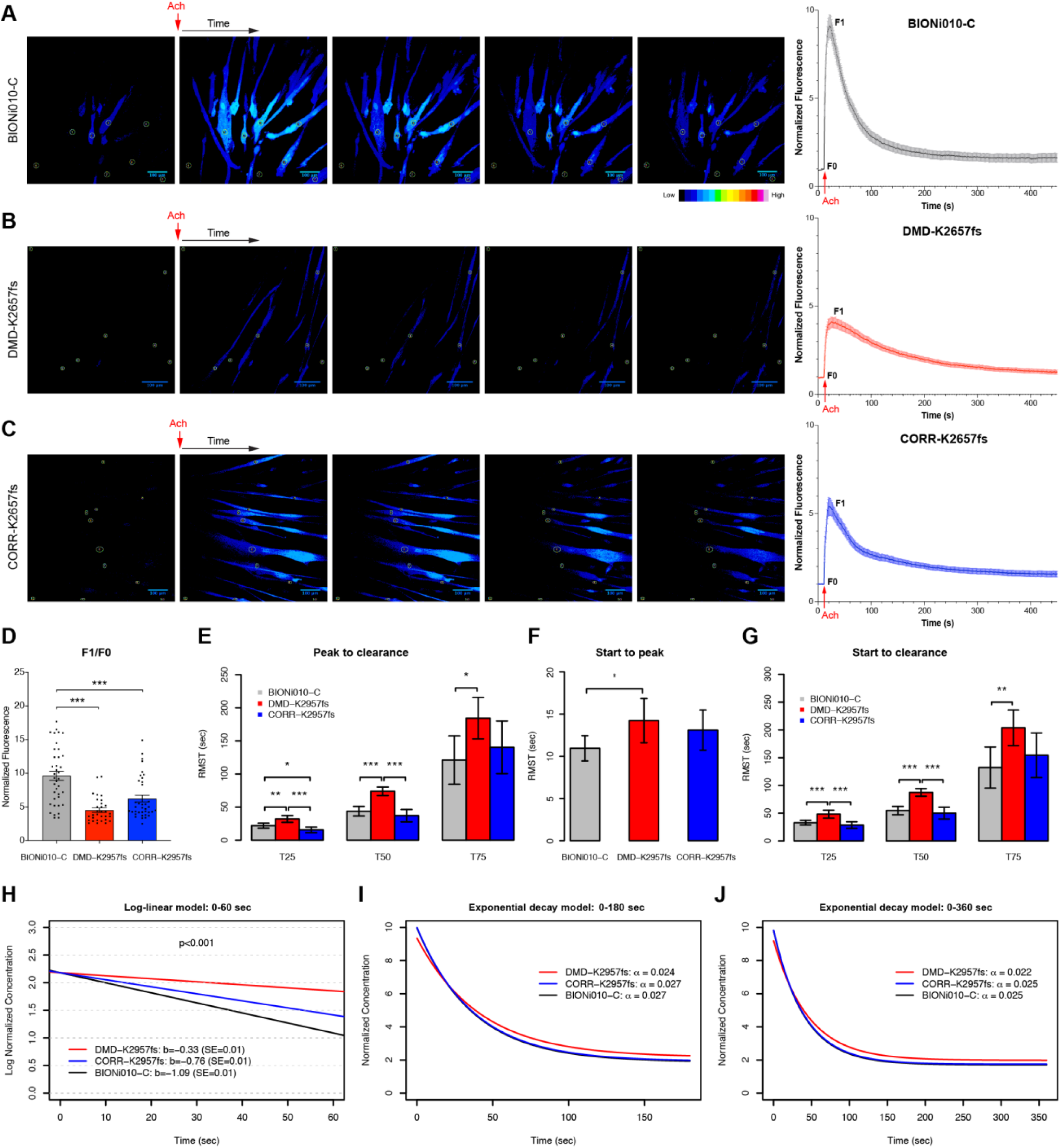
Analysis of intracellular Ca^2+^ transients and mathematical modeling. (A-C) Representative images of intracellular Ca^2+^ flux stimulated by acetylcholine (Ach) in BIONi010-C (A, n= 33), DMD-K2957fs (B, n=31), and CORR-K2957fs (C, n= 35) myotubes over time. Color codes indicate the levels of fluorescent signal. Scale bar, 100 μm. Data were collected from at least three independent experiments on Day 5 of secondary myogenic differentiation cultures. Circle areas indicate regions of interest (ROI). The kinetics of Ca^2+^ transients of each group was plotted, respectively. Data are mean ± SEM. F0, baseline of fluorescence. F1, peak of fluorescence. (D) The analysis of Ca^2+^ transient kinetics shows that BIONi010-C has significantly higher mean value of F1/F0 than DMD-K2957fs and CORR-K2957fs myotubes. Data are mean ± SEM. One-way ANOVA, Tukey test (*** p < 0.001). (E) Restricted mean time from peak to 25, 50 and 75 percent clearance. Data are estimated mean ± 95%CI, Z-test (* p < 0.05, ** p < 0.01, *** p < 0.001). (F) Restricted mean time from start to peak. Data are estimated mean ± 95%CI, Z-test (* p < 0.05). (G) Restricted mean time from start to 25, 50 and 75 percent clearance. Data are estimated mean ± 95%CI, Z-test (* p < 0.05, ** p < 0.01, *** p < 0.001). (H) Log-linear model for elimination: 0-60 seconds. (I) Exponential decay model for clearance: 0-180 seconds. (J) Exponential decay model for clearance: 0-360 seconds. (I and J) Note that the Ca^2+^ clearance rates in the CORR-K2957fs (blue line) and BIONi010-C (black line) myotubes are completely overlapping.

Next, we investigated the dynamics of the return of the Ca^2+^ transient to baseline which we termed intracellular Ca^2+^ clearance. Time to 25%, 50% and 75% reduction to baseline values are naturally skewed to the right and subject to censoring. Therefore, analysis of these data based strictly on observed values may result in bias due to misspecification of the data distribution and missing data values. Both issues are readily addressed within survival analysis framework widely used in clinical trials and observational studies. We utilized here the Restricted Mean Survival Time (RMST) approach, that provides absolute measure of time to the event of interest [39]. The RMST is defined as the area under the curve of the survival function up to a time *τ* < ∞.

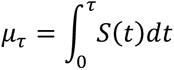

where *S*(*t*) can be estimated by 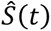, the Kaplan-Meier estimator, and *τ* is set to the minimum of the largest observed time in each of the two groups under comparison. The RMST is the population average of the time before event of interest during first *τ* seconds of follow-up.

Our analysis showed that DMD-K2957fs myotubes had significantly longer time from peak to 25%, 50% and 75% Ca^2+^ clearance (T25, T50 and T75) compared to BIONi010-C, and significantly longer time to 25% and 50% clearance compared to CORR-K2957fs. In addition, BIONi010-C myotubes had significantly longer time from peak to 25% Ca^2+^ clearance than CORR-K2957fs (Fig. 4E; Table S4). Further analysis of the restricted mean time from addition of acetylcholine to peak of Ca^2+^ transients showed that DMD-K2957fs myotubes had significantly longer time from start of experiment to peak (T-peak) compared to BIONi010-C, but no significant difference compared to CORR-K2957fs (Fig. 4F; Table S4). Finally, we investigated the restricted mean time from start of experiment to 25%, 50% and 75% Ca^2+^ clearance. DMD-K2957fs myotubes had significantly longer time from start to 25%, 50% and 75% clearance compared to BIONi010-C, and significantly longer time from start to 25% and 50% clearance compared to CORR-K2957fs (Fig. 4G; Table S4). Taken together, these results suggest slowing of intracellular cytoplasmic Ca^2+^ removal in DMD-K2957fs myotubes, which may contribute to alterations in E-C coupling.

### 3.6 Mathematical modeling reveals reduced intracellular Ca^2+^ clearance rate in DMD-K2957fs myotubes

To further elucidate the defective intracellular Ca^2+^ clearance in DMD-K2957fs myotubes, we sought to simulate the Ca^2+^ clearance rate using log-linear and exponential decay models.

#### Log-linear model for elimination

The elimination rate constant *k_e_* is defined as follows.

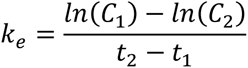

where *t_i_* is time, *C_i_* is concentration at time *i*. The elimination constant can be viewed as a linear slope *b* for concentration change on the log scale (Fig. 4H). The mixed effect repeated measures model was used to estimate linear trends and to perform comparison on the log concentration scale. The model included fixed effects of time, group and time by group interaction. The intercept was treated as random. Variance covariance structure of compound symmetry form was assumed between observations for each sample. The log-linear model (0-60 seconds) showed that DMD-K2957fs myotubes had the shallowest slope (-0.33 log-concentration/min), compared with CORR-K2957fs (- 0.76 log-concentration/min) and BIONi010-C myotubes (-1.09 log-concentration/min), suggesting the intracellular Ca^2+^ elimination rate in DMD-K2957fs myotubes is significantly slower than that in CORR-K2957fs or BIONi010-C myotubes(Fig. 4H).

#### Exponential decay model for clearance

Next, we fit the exponential decay models to the normalized concentration on interval of 0-180 seconds past peak concentration. The peak concentration time is sample specific. The exponential decay model is specified as follows.

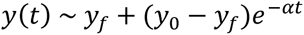

where the measured value *y* starts at *y*_0_ and decays towards *y_f_* at a rate *α* (Fig. 4I; Table S5). The exponential decay model (0-180 seconds) showed that DMD-K2957fs myotubes had slower Ca^2+^ decay rate (0.024 sec^-1^), compared with CORR-K2957fs (0.027 sec^-1^) or BIONi010-C myotubes (0.027 sec^-1^). Similarly, we fit the exponential decay models to the normalized concentration on interval of 0- 360 seconds past peak concentration. The peak concentration time is sample specific (Fig. 4J; Table S5). The exponential decay model (0-360 seconds) showed that DMD-K2957fs myotubes had slower Ca^2+^ decay rate (0.022 sec^-1^) than CORR-K2957fs and BIONi010-C myotubes, which had comparable Ca^2+^ decay rate (0.025 sec^-1^). Taken together, both the log-linear elimination and exponential decay models suggest that DMD-K2957fs myotubes had significantly slower clearance rate of intracellular Ca^2+^ than CORR-K2957fs and BIONi010-C myotubes.

### 3.7 Cytoplasmic Ca^2+^ flux in myotubes is determined by intracellular and extracellular sources

We next examined the pharmacological dependence of the Ca^2+^ transient on intracellular and extracellular sources of Ca^2+^. Prior to acetylcholine stimulation, we used ryanodine to block ryanodine receptors, cyclopiazonic acid to block SERCA, or nifedipine to block L-type voltage-dependent Ca^2+^ channels (Ca_V_1.1) in BIONi010-C, DMD-K2957fs, and CORR-K2957fs myotubes (Fig. 5A).

**Fig. 5.**
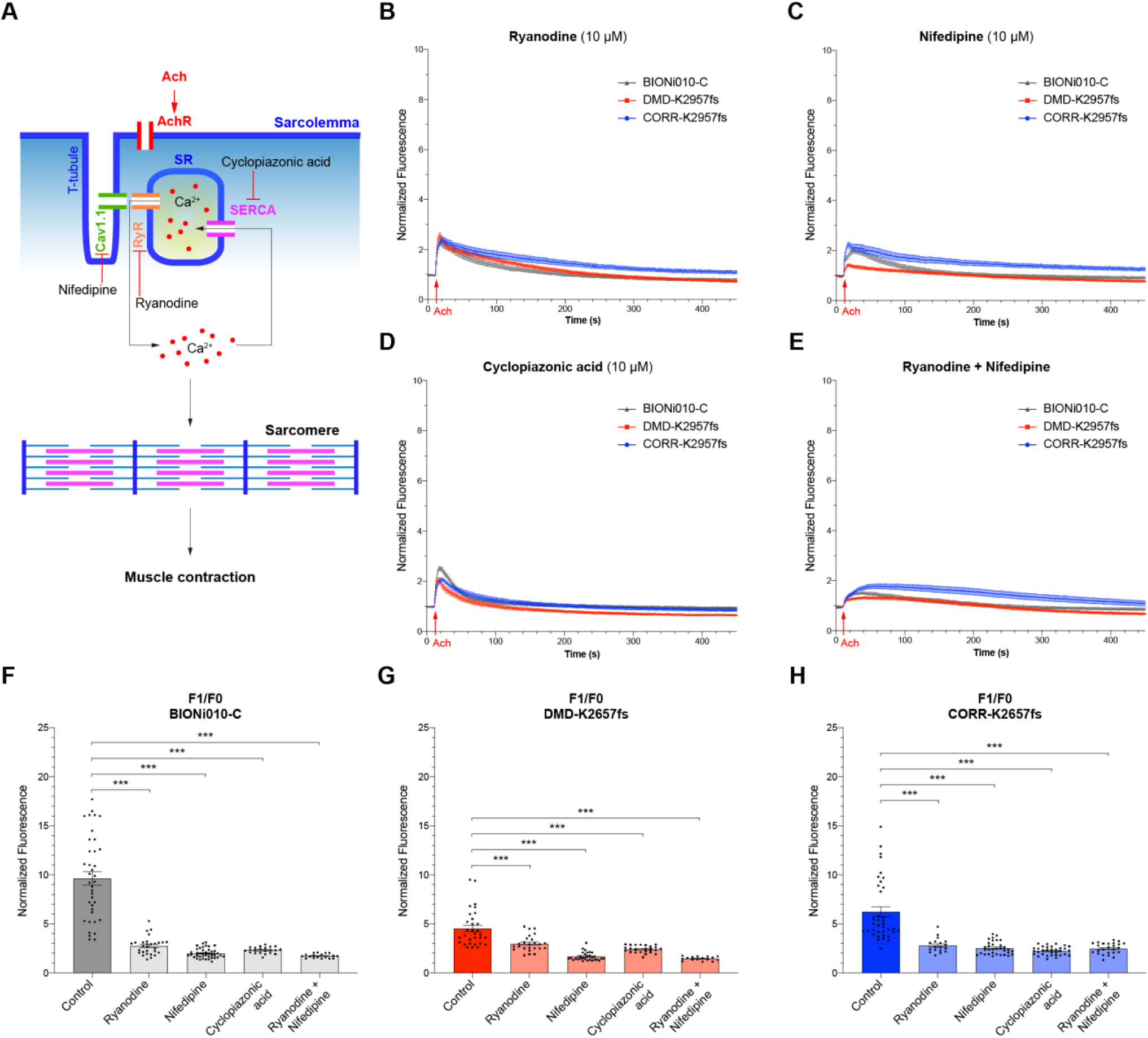
Analysis of calcium channel blockers on acetylcholine-stimulated intracellular Ca^2+^ transients. (A) A schema of E-C coupling and calcium channel blockers. Acetylcholine (Ach) binds to acetylcholine receptors (AchR) on the sarcolemma of muscle fibers, leading to the depolarization of membrane, which spreads to the T-tubules, causing a conformational change of Ca_v_1.1, which in turn activates the opening of RYR to release Ca^2+^ from the SR lumen into the cytoplasm. This leads to activation of actin-myosin sliding within the sarcomere, resulting in muscle contraction. During the relaxation of skeletal muscle, SERCA transfers cytosolic Ca^2+^ back to the lumen of SR. Calcium channel blockers: ryanodine blocks RYR, nifedipine blocks Ca_v_1.1, and cyclopiazonic acid blocks SERCA. (B-E) The kinetics of Ca^2+^ transient plots of Day 5 secondary myogenic differentiation cultures incubated with ryanodine (B; BIONi010-C, n= 29; DMD-K2957fs, n= 25; CORR-K2957fs, n= 18), nifedipine (C; BIONi010-C, n= 41; DMD-K2957fs, n= 30; CORR-K2957fs, n= 33), cyclopiazonic acid (D; BIONi010-C, n= 22; DMD-K2957fs, n= 24; CORR-K2957fs, n= 33) or ryanodine plus nifedipine (E; BIONi010-C, n= 18; DMD-K2957fs, n= 16; CORR-K2957fs, n= 25), prior to acetylcholine stimulation. Data represent mean ± SEM (at least three independent experiments). F0, baseline of fluorescence. F1, peak of fluorescence. (F-H) Compared to untreated controls (data from Fig. 4D), the mean Ca^2+^ transient amplitudes (F1/F0) in myotubes treated with calcium channel blockers were all significantly reduced. BIONi010-C (F); DMD-K2957fs (G); CORR-K2957fs (H). Data are mean ± SEM. One-way ANOVA, Tukey test (*** p < 0.001).

Incubation with ryanodine, nifedipine or cyclopiazonic acid all reduced the magnitude of the Ca^2+^ transients in either BIONi010-C, DMD-K2957fs or CORR-K2957fs myotubes (Fig. 5B-D), suggesting a dependence on release from the SR and entry via the sarcolemma through the L-type calcium channel. The co-application of ryanodine and nifedipine resulted in slower increase in cytosolic Ca^2+^ different from the rapid calcium transient seen in other circumstances (Fig. 5E). Compared to untreated controls, quantification of Ca^2+^ transient amplitude in myotubes treated with calcium channel blockers showed a significant reduction of F1/F0 in either BIONi010-C, DMD-K2957fs or CORR-K2957fs myotubes (Fig. 5F-H). While Ca^2+^ entry through Ca_V_1.1 is not necessary for E-C coupling in mammalian skeletal muscle [40], our results suggest that Ca^2+^ flux in the human PSC-derived myotubes was determined by both intracellular and extracellular sources. In addition, the fact that Ca^2+^ transients in human PSC-derived myotubes responded to pharmacological treatments suggest that our model should be amenable to drug testing for identifying compounds that can ameliorate DMD phenotypes.

### 3.8 Velocity of contractility is significantly reduced in DMD-K2957fs myotubes

Apart from genes involved in Ca^2+^ handling, GSEA also identified several sarcomere components within the leading-edge subset of the MYOGENESIS gene set, including troponin (*TNNC2*, *TNNI1*, *TNNI2*, *TNNT1*, *TNNT3*) and tropomyosin (*TPM2*), which were significantly down-regulated in DMD- K2957fs compared to CORR-K2957fs myogenic cultures (Fig. 6A). Real-time qPCR confirmed similar trends of differential gene expression between DMD-K2957fs and CORR-K2957fs with statistical significance in *TNNI2* and *TPM2* (Fig. S3B). As troponin and tropomyosin regulate muscle contraction through Ca^2+^ binding [41], we sought to determine whether loss of dystrophin may affect myotube contractility. To do this, we measured the velocity of myotube contraction in response to acetylcholine stimulation using particle image velocimetry (PIV) tool in MATLAB, PIVlab [24]. The algorithm of PIVlab computes the displacement of a field of pixels between two consecutive images, frame by frame, within a video recording. PIVlab analysis showed that DMD-K2957fs myotubes had significantly lower maximum velocity after acetylcholine stimulation, compared to BIONi010-C or CORR-K2957fs myotubes (Fig. 6B,C). The release of Ca^2+^ from the SR into the cytosol activates actin-myosin sliding within the sarcomere that leads to muscle contraction (Fig. 5A). Despite significantly higher Ca^2+^ transient amplitude (F1/F0) in BIONi010-C myotubes than CORR-K2957fs or DMD-K2957fs myotubes (Fig. 4D), there was no significant difference between maximum velocities of BIONi010-C and CORR-K2957fs myotube contractility (Fig. 6D). Together, our findings suggest that down-regulated gene expression of troponin and tropomyosin in the absence of dystrophin may contribute to the reduced velocity of myotube contractility.

**Fig. 6.**
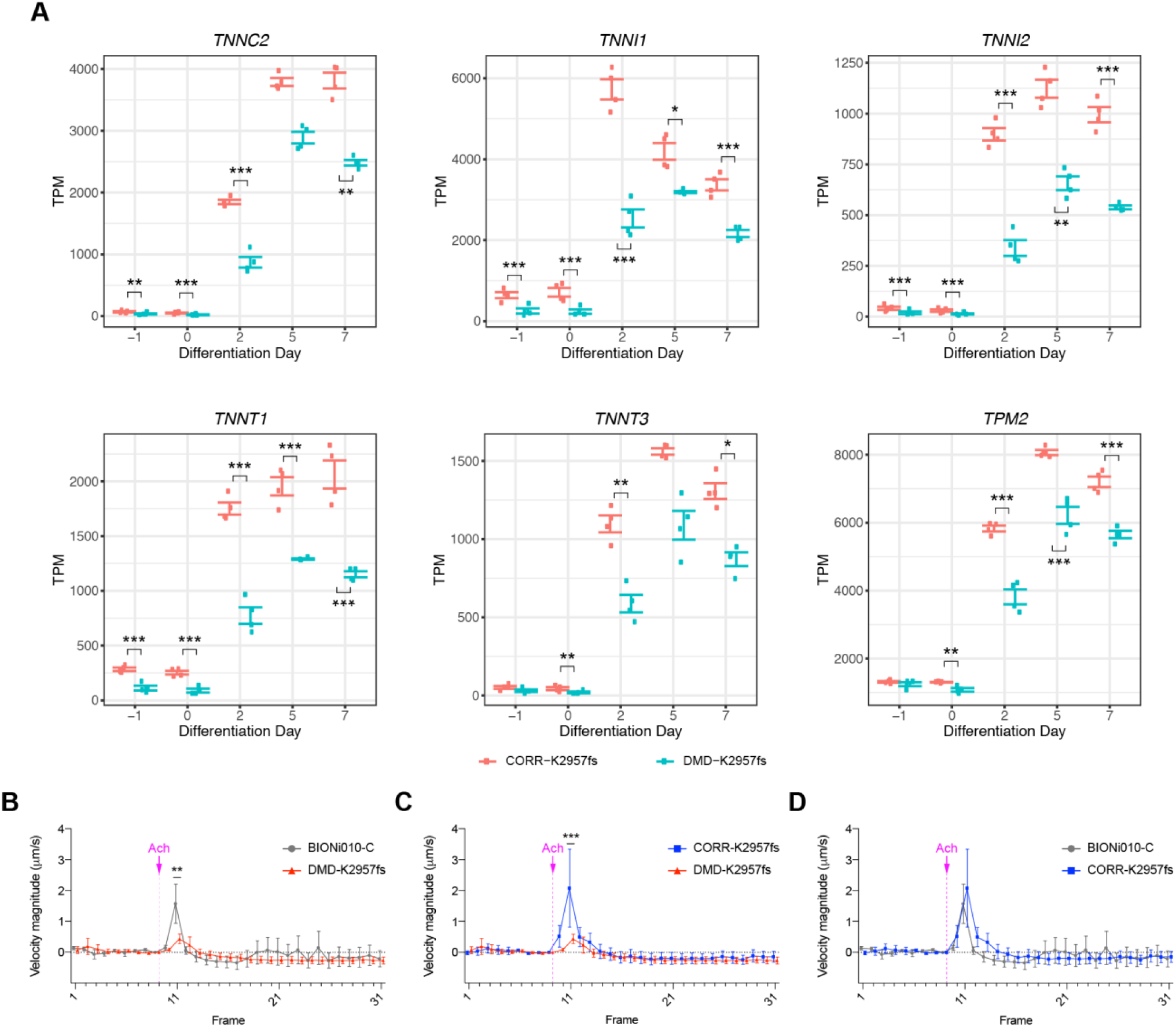
Analysis of muscle contraction genes and myotube contractility. (A) Time-course expression levels of genes within HALLMARK_MYOGENESIS that regulate muscle contraction, including *TNNC2*, *TNNI1*, *TNNI2*, *TNNT1*, *TNNT3* and *TPM2*. Wald test with Benjamini-Hochberg method for multiple comparison adjustment (* padj < 0.05, ** padj < 0.01, *** padj < 0.001). TPM, Transcripts Per Million. Each condition has 4 biological replicates. (B-D) Analysis of myotube contraction velocity using Particle Image Velocimetry. In response to acetylcholine (Ach) stimulation, DMD-K2957fs myotubes had significantly slower velocity magnitude than BIONi010-C (B) or CORR-K2957fs (C) myotubes, whereas no significant difference of velocity between BIONi010-C and CORR-K2957fs myotubes (D). Data represent mean ± SEM (n= 9; at least three independent experiments). Two-way ANOVA, Sidak test (** p < 0.01; *** p < 0.001).

## 4. Discussion

Our study establishes a new DMD patient-derived PSC model of skeletal muscle carrying a *DMD* c.8868delC mutation in exon 59 (DMD-K2957fs), resulting in the absence of full-length dystrophin isoform (Dp427). Precise correction of the *DMD* c.8868delC mutation using CRISPR-mediated genome editing generates an isogenic control (CORR-K2957fs), which restores the expression of full-length dystrophin. Interestingly, GSEA comparing DMD-K2957fs to CORR-K2957fs transcriptomes during secondary myogenic differentiation identifies affected HALLMARK gene sets that regulate inflammation related pathways, fibrosis and myogenesis. In particular, our analysis shows that HALLMARK gene sets INTERFERON GAMMA RESPONSE and TNFA SIGNALING VIA NFKB are significantly enriched in DMD transcriptomes (Day 2, 5 and 7). In agreement with our findings, studies have shown that interferon-γ and TNF-induced activation of NF-κB are involved in suppressing muscle-specific gene expression (e.g. *MyoD* mRNA) and in skeletal muscle wasting [42, 43]. Genetic ablation of myofiber IKKβ/NF-κB signaling in *mdx* mice promotes muscle regeneration [44]. In addition, we show that the IL6 JAK STAT3 SIGNALING and TGF BETA SIGNALING gene sets are enriched in DMD transcriptomes. Consistent with our results, inhibition of IL6/JAK/STAT signaling stimulates muscle regeneration and ameliorates dystrophic phenotypes in *mdx* mice [45, 46]. Furthermore, increased TGFβ1 expression correlates with fibrosis in DMD patients [47]. Taken together, our results suggest that DMD myogenic transcriptomes reflect aberrant signaling pathways and our cell model may have a predisposition towards pathophysiology described in DMD patients.

In agreement with our transcriptome analysis, a recent multi-omic study by Mournetas *et al*. [48] shows that human PSC-derived myotubes resemble fetal-like phenotypes and are associated with several aspects of DMD defects, including Ca^2+^ homeostasis (e.g. *ATP2A2* mRNA; RYR1 and CACNA1S proteins) and markers of fibrosis. Contrast to our study and others [35], up-regulation of TGFβ signaling in DMD was not reported in the study by Mournetas *et al*., which may be due to some fundamental differences between differentiation protocols and media [22, 49], as well as inter-individual variability. The transcriptome analysis by Mournetas *et al*. was carried out from human PSCs (Day 0) to myotubes (Day 25) and the cell culture media contain specific molecules that inhibit TGFβ signaling from Day 0 to Day 17 of myogenic differentiation. In contrast, our transcriptome analysis was carried out from human PSC-derived MPCs (Day -1 and 0) to myotubes (Day 7) and our differentiation medium did not contain any inhibitors for TGFβ signaling. An interesting observation in our present study is the reduced percentage of PAX7-positive cells in DMD-K2957fs myogenic cultures consistent with a recent study modeling the DMD-R3381X mutation affecting all dystrophin isoforms [36]. While significantly down-regulated *PAX7* transcripts in DMD-K2957fs myogenic transcriptomes correlate with the reduced percentage of PAX7-positive cells, it should be noted that *PAX7* expression is not significantly down-regulated in DMD-R3381X myogenic transcriptomes, suggesting that *PAX7* translation or other mechanisms may be involved. We reason that differences of *PAX7* transcript levels in the two DMD myogenic transcriptomes could be due to mRNA stability and/or alterations in epigenetic regulatory processes. Altogether, further comparisons between studies of DMD myogenic transcriptomes with experimental validation will shed light on identifying common pathological mechanisms in dystrophin-deficient skeletal muscle.

In contrast to gene sets enriched in DMD-K2957fs, HALLMARK MYOGENESIS is enriched in CORR-K2957fs during secondary myogenic differentiation. Investigating the leading-edge subset of the MYOGENESIS gene set reveals that several core enrichment genes involved in Ca^2+^ handling are significantly down-regulated in DMD-K2957fs transcriptomes, such as *RYR1*, *ATP2A1*, *SLN*, *CASQ2*, and *CACNA1H*, as well as *CACNA1S* (Ca_V_1.1), suggesting dysregulation of Ca^2+^ homeostasis and E-C coupling in DMD. In support of our findings, previous studies showed that calsequestrin is reduced in *mdx* mouse skeletal muscles using subproteomics analysis [50] and that measurement of RYR permeability reveals a role of calsequestrin in termination of SR Ca^2+^ release in skeletal muscle [51].

Moreover, *CACNA1H* downregulation induces skeletal muscle atrophy *in vitro* and in *Cacna1h−/−* mice [52]. Interestingly, Woods *et al*. show that the amplitude of the intracellular Ca^2+^ increase caused by electrical stimulation is impaired in *mdx* mouse muscle fibers [53]. In our study using acetylcholine stimulation, we also observe reduced intracellular Ca^2+^ amplitude (F1/F0) in DMD-K2957fs myotubes, compared to BIONi010-C and CORR-K2957fs myotubes. The difference of F1/F0 between BIONi010-C and CORR-K2957fs myotubes may reflect inter-individual variability. Based on our transcriptome analysis, we reason that the reduced SR Ca^2+^ release flux may be due to a reduction of RYR opening and decreased calsequestrin in the SR [51]. Alternatively, but not mutually exclusive, it may be explained by reduced Ca^2+^ storage and buffering in the SR [50]. Furthermore, aberrantly down- regulated *ATP2A1* expression in DMD-K2957fs myogenic cultures suggest that cytosolic Ca^2+^ reuptake may be affected in DMD skeletal muscle upon acetylcholine stimulated SR Ca^2+^ release and prompt us to investigate the Ca^2+^ transient kinetics. Importantly, our RMST analysis and mathematical modeling demonstrate that DMD patient PSC-derived myotubes have significantly reduced intracellular Ca^2+^ clearance rate, compared to the CRISPR-corrected isogenic control and non-isogenic wildtype myotubes. Consistent with our findings, Woods *et al*. also observed that *mdx* mouse muscle fibers have a marked prolongation of the intracellular Ca^2+^ decay [53]. Regarding the sources of Ca^2+^ transients, our pharmacological analysis indicates that Ca^2+^ flux in human PSC-derived myotubes is determined by intracellular and extracellular sources. While compelling evidence suggests that DMD patient PSC-derived myotubes are associated with abnormal Ca^2+^ handling and alternation of E-C coupling specifically slows removal of cytoplasmic Ca^2+^, how loss of dystrophin leads to dysregulation of Ca^2+^ homeostasis in skeletal muscle will require further investigation. Furthermore, this prolonged Ca^2+^ transient may have effects on gene transcription and in part account for the changes in transcriptional profile [54–56].

In addition to Ca^2+^ handling genes, core enrichment genes in the MYOGENESIS gene set also include *TNNC2, TNNI1, TNNI2, TNNT1, TNNT3* and *TPM2*, which are down-regulated in DMD-K2957fs transcriptomes. These genes encode troponin complex components and tropomyosin 2, which regulate muscle contraction through Ca^2+^ binding [41]. Consistent with the transcriptome analysis, we show that the maximum velocity of myotube contractility is significantly reduced in DMD-K2957fs, compared to the isogenic control CORR-K2957fs or non-isogenic wildtype BIONi010-C. Taken together, our results reinforce the notion that Ca^2+^ signaling and homeostasis play important roles in modulating skeletal muscle function and may underlie the pathogenesis of DMD.

Elevated intracellular Ca^2+^ has been associated with increased cellular stress conditions, such as the production of reactive oxygen species (ROS) and mitochondrial damage. ER stress and unfolded protein response have been reported in animal models of muscular dystrophies due to mutations affecting the DGC function [57, 58]. It is worth noting that our GSEA results show that HALLMARK gene sets REACTIVE OXYGEN SPECIES PATHWAY and UNFOLDED PROTEIN RESPONSE are enriched in DMD transcriptomes. Interestingly, it has been reported that SERCA activity declines progressively in *mdx* mouse muscle under conditions of cellular stress [59]. Together, our results suggest that reduced intracellular Ca^2+^ reuptake rate in DMD muscle fibers may lead to greater mitochondrial damage and dysfunction [60], resulting in a vicious cycle of cellular stress, changes of gene expression, activation of proteases (e.g. calpain) and eventual muscle cell death. In support of our results, studies have shown that overexpression of SERCA can increase SERCA activity and ameliorate histopathological and biochemical features of muscular dystrophy in the *mdx* (mild) and *mdx:utr* (severe) mouse models [61, 62] and that modulation of SERCA function by increasing heat shock protein 72 (Hsp72) preserves muscle function and improves the dystrophic pathology in the *mdx* and *mdx:utr* mice [59]. Finally, adeno-associated virus-mediated SERCA2a gene therapy in *mdx* mice enhanced Ca^2+^ uptake in the skeletal and cardiac muscle, improved muscle strength and ameliorated dilated cardiomyopathy [63].

It has been reported that sarcolipin protein expression is abnormally up-regulated in the *mdx* and *mdx:utr* dystrophic mouse models, while SERCA1 protein expression is reduced, and that reducing sarcolipin expression mitigates skeletal muscle and cardiac pathology in the *mdx:utr* mice [64]. Although reduced SERCA1 in *mdx* and *mdx:utr* mice is consistent with the down-regulated *ATP2A1* in our transcriptome analysis, we did not observe significantly up-regulated sarcolipin (*SLN*) gene expression. In fact, *SLN* expression was down-regulated in the DMD-K2957fs myogenic cultures. We speculate this discrepancy may reflect the differences of transcriptional regulation between rodent and human species. This reiterates the importance of using human-relevant models that recapitulate human pathophysiology for studying disease mechanisms.

Our study provides new insights into the physiological and pharmacological properties of Ca^2+^ in skeletal muscle generated from DMD patient-derived PSCs however it has limitations. It will be important in future work to dissect E-C coupling in more detail and the maturity of the response compared to native adult skeletal muscle. More work has been undertaken in cardiac muscle and it is recognized that many standard differentiation protocols lead to an immature electrophysiological and contractile phenotype [65]. For example, it appears in our cells that the Ca^2+^ transient is also dependent on extracellular calcium entry which is inconsistent with a large dependence of the transient on SR Ca^2+^ release mediated by a direct molecular interaction between the L-type Ca^2+^ channel and ryanodine receptor. We also assume that the initial events after acetylcholine binding to nicotinic acetylcholine receptors is membrane depolarization and action potential generation but we do not have direct electrophysiological measurements of this. Finally, Fluo-4 is a non-ratiometric dye and we did not titrate the dye to resting Ca^2+^ concentration. These areas should be addressed by more systematic and detailed studies in the future.

Since dilated cardiomyopathy is now a leading cause of death among DMD/BMD patients, it will be of interest to investigate mechanisms of Ca^2+^ dysregulation underlying DMD-PSC derived cardiomyocytes. In agreement with our findings, Kyrychenko *et al.* show that DMD patient iPSC-derived cardiomyocytes have significantly slower intracellular Ca^2+^ reuptake, compared to isogenic controls generated by CRISPR-mediated exon deletion strategies that restored the dystrophin reading frame [66]. It should be noted that strategies utilizing in-frame deletions of the *DMD* mutations may not fully restore dystrophin function as reflected by the varying degrees of iPSC-cardiomyocyte function after gene editing [66]. This highlights the importance of pre-clinical assessments when designing mini- and micro-dystrophin gene therapies.

Our study has limitations in its representation of DMD pathophysiology because only one patient-derived PSC line (DMD-K2957fs) and one CRISPR-corrected line (CORR-K2957fs) were used without analysis in clonal lines. While our results show that the CRISPR-corrected myotubes (CORR- K2957fs) physiologically resemble the non-isogenic wildtype myotubes (BIONi010-C) in terms of intracellular Ca^2+^ clearance rate and myotube contractility, future studies employing multiple DMD patient-derived lines with isogenic controls and analysis in clonal lines will address the range of variability of these measures once a larger number of PSC clones are evaluated.

Each year, around 20,000 children worldwide are born with DMD. Overall, it is estimated that there are about 300,000 DMD patients. A significant economic burden is associated with DMD and increases markedly with disease progression [67]. Current standards of care for DMD patients are corticosteroids prednisone and deflazacort [68]. Despite improved muscle function and delay disease progression, it should be noted that long term administration of steroids has been associated with several side effects, such as excessive weight gain, delayed growth and puberty, increased risk of osteoporosis and behavioral issues. Currently, the only approved treatments designed to target patients with mutations that cause DMD are Translarna, EXONDYS 51, VYONDYS 53/Viltepso and AMONDYS 45, cumulatively suitable for treating up to ∼30% of DMD patients. Thus, it is important to develop effective treatments for the remaining DMD patients.

In summary, our study suggests that Ca^2+^ handling pathways amenable to pharmacological modulation are potential therapeutic targets for DMD. Importantly, our work provides an isogenic human physiology-relevant platform for elucidating mechanisms underlying DMD and rapid pre-clinical assessment of therapeutic strategies, such as mini-/micro-dystrophin gene therapies.

## Acknowledgements

We thank the assistance from Dr Luke Gammon at the Blizard Phenotypic Screening Facility. The support of the MRC Centre for Neuromuscular Diseases Biobank and Pierpaolo Ala are gratefully acknowledged. This work was facilitated by the NIHR Cardiovascular Biomedical Research Centre at Barts. This work was funded by a Pfizer research grant to YYL and AT. JEM and JM are funded by Muscular Dystrophy UK (grant 17GRO-PG36-0165).

## Supplementary Material

**Fig. S1.**
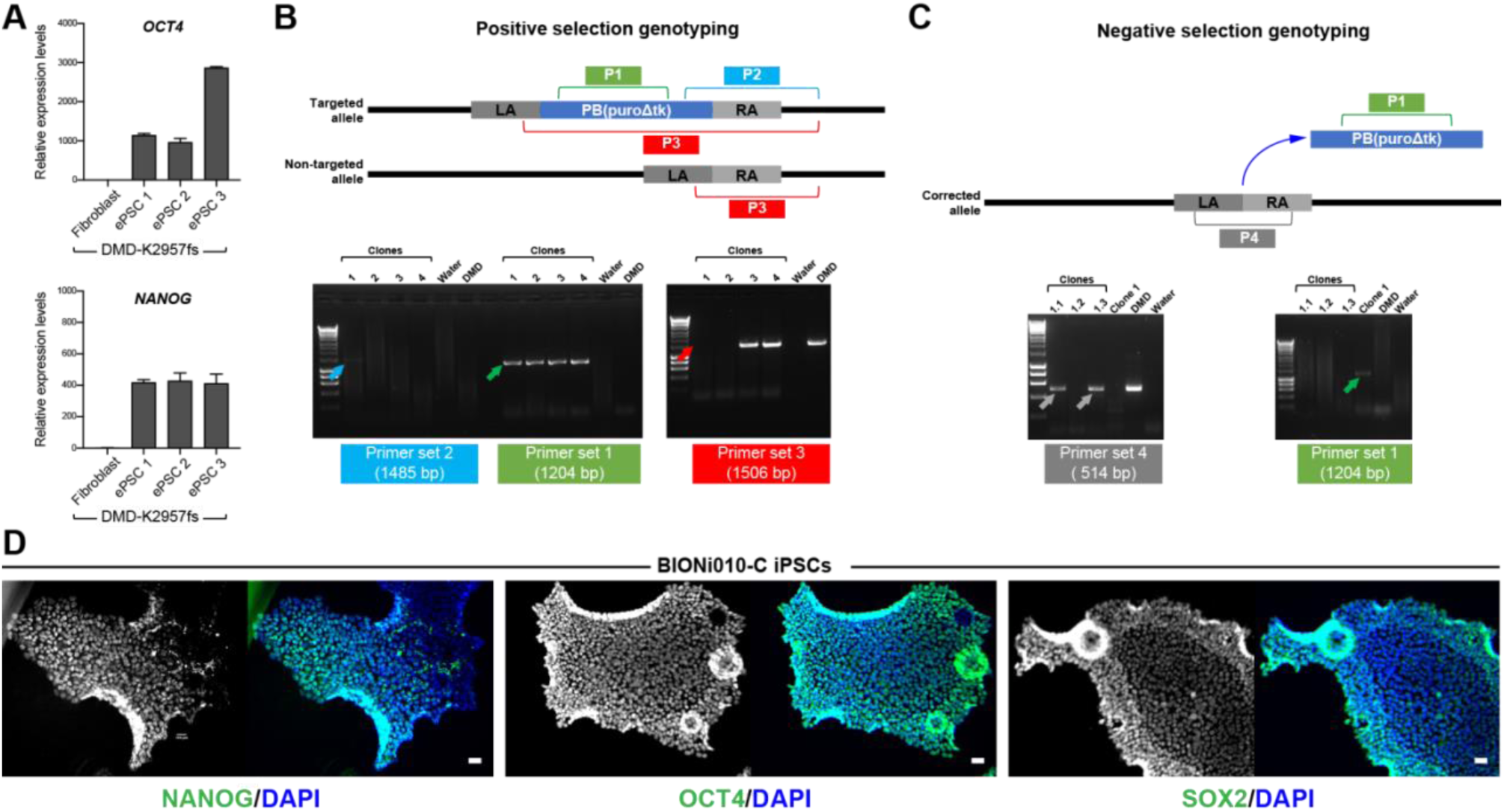
Characterization of human PSC lines and genotyping of genome editing. (A) Gene expression analysis of DMD-K2957fs ePSC clonal lines showed high levels of *OCT4* and *NANOG* mRNA expression, compared with parental fibroblasts. Values are mean ± SD (n=3). (B) PCR genotyping to identify targeted clones after puromycin positive selection. Successfully target clones were positive for PCR product 1 (green) and 2 (blue), while negative for PCR product 3 (red), e.g. clone 1. LA, left homology arm; RA, right homology arm. (C) PCR genotyping to identify modified clones after FIAU negative selection. Successfully modified clones were positive for PCR product 4 (grey), while negative for PCR product 1 (green), e.g. clone 1.1 and 1.3. (D) The BIONi010-C iPSC line expressed characteristic pluripotency markers, NANOG, OCT4 and SOX2. Scale bar, 100 μm.’

**Fig. S2.**
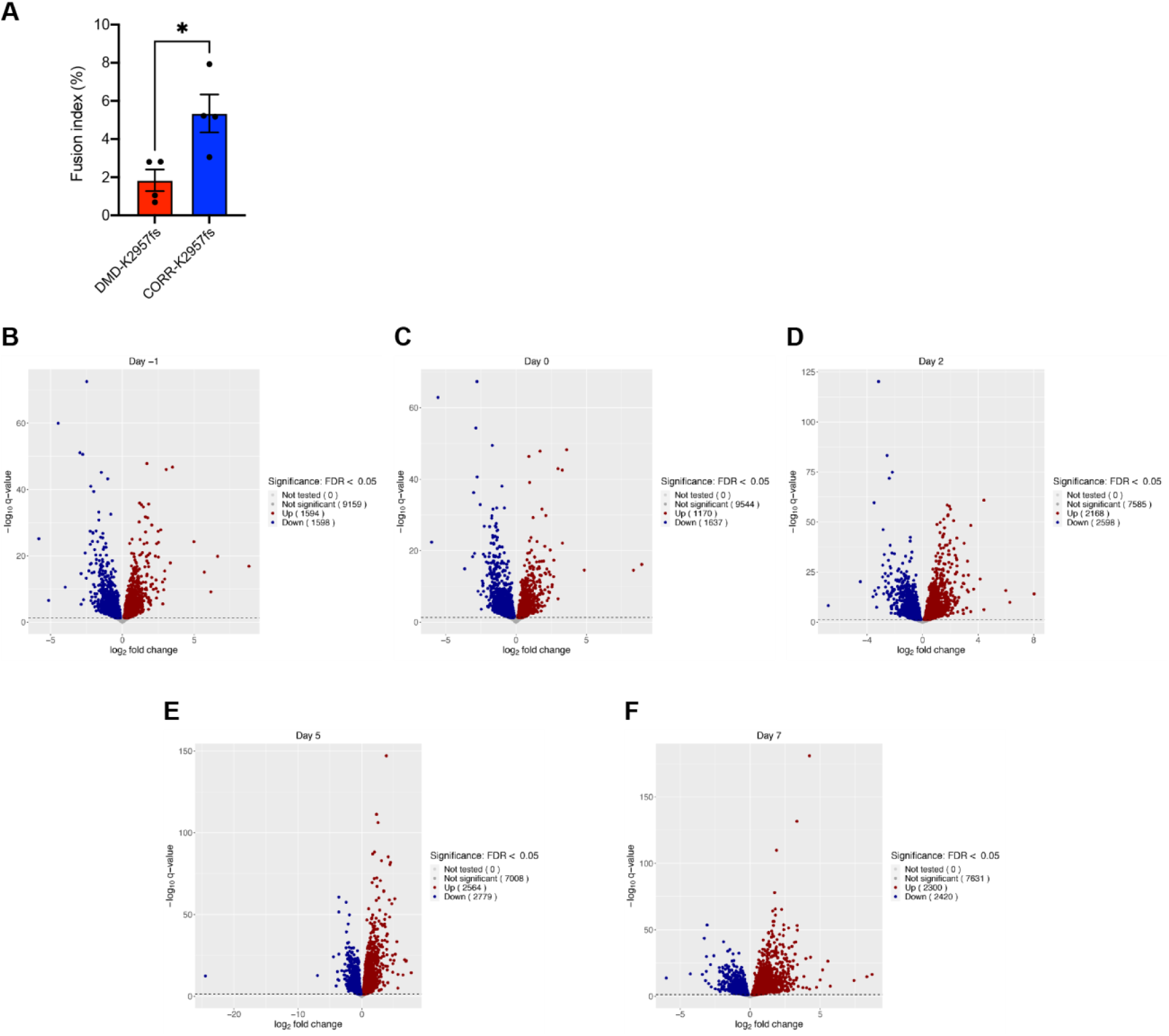
(A) Fusion index of DMD-K2957fs and CORR-K2957fs myogenic cultures at Day 5. Fusion index was defined the number of nuclei within myotubes (≥ 3 nuclei) as a percentage of total nuclei. Data are mean ± SEM (n= 55; 4 independent experiments). Unpaired t-test (* p < 0.05). (B-F) Differential gene expression between DMD-K2957fs and CORR-K2957fs transcriptomes at Day -1, 0, 2, 5 and 7. The number of significantly up-regulated (red) and down-regulated (blue) genes in each time point are specified (FDR < 0.05).

**Fig. S3.**
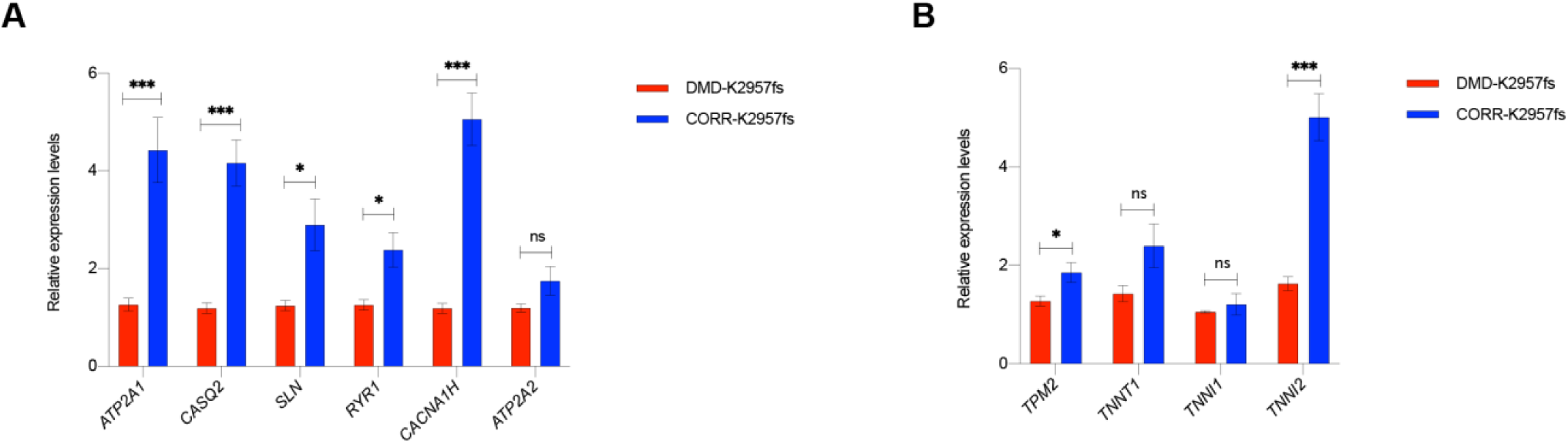
Real-time qPCR analysis of Ca^2+^ handling and muscle contraction genes in DMD-K2957fs and CORR-K2957fs myogenic cultures. (A) Relative gene expression levels of *ATP2A1*, *CASQ2*, *SLN*, *RYR1*, *CACNA1H* and *ATP2A2* in DMD-K2957fs and CORR-K2957fs myogenic cultures (Day 5). (B) Relative gene expression levels of *TPM2*, *TNNT1*, *TNNI1*, and *TNNI2* in DMD-K2957fs and CORR-K2957fs myogenic cultures (Day 5). Data represent mean ± SEM (n= 6 to 9; three independent experiments). Unpaired t-test (ns, not significant; * p < 0.05; *** p < 0.001).

**Table S1.**
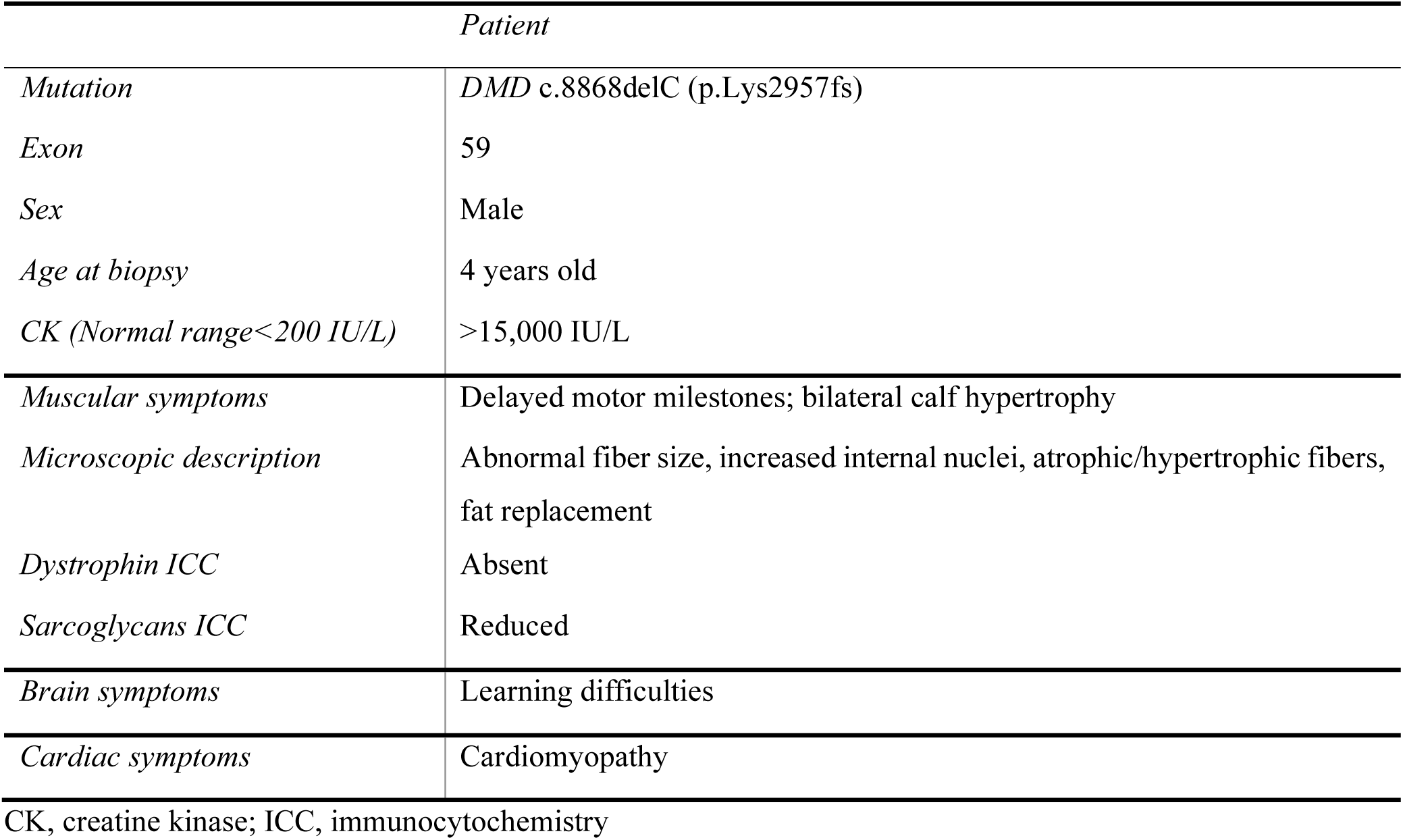
Summary of the DMD patient’s mutation, muscle biopsy report and symptoms

**Table S2.**
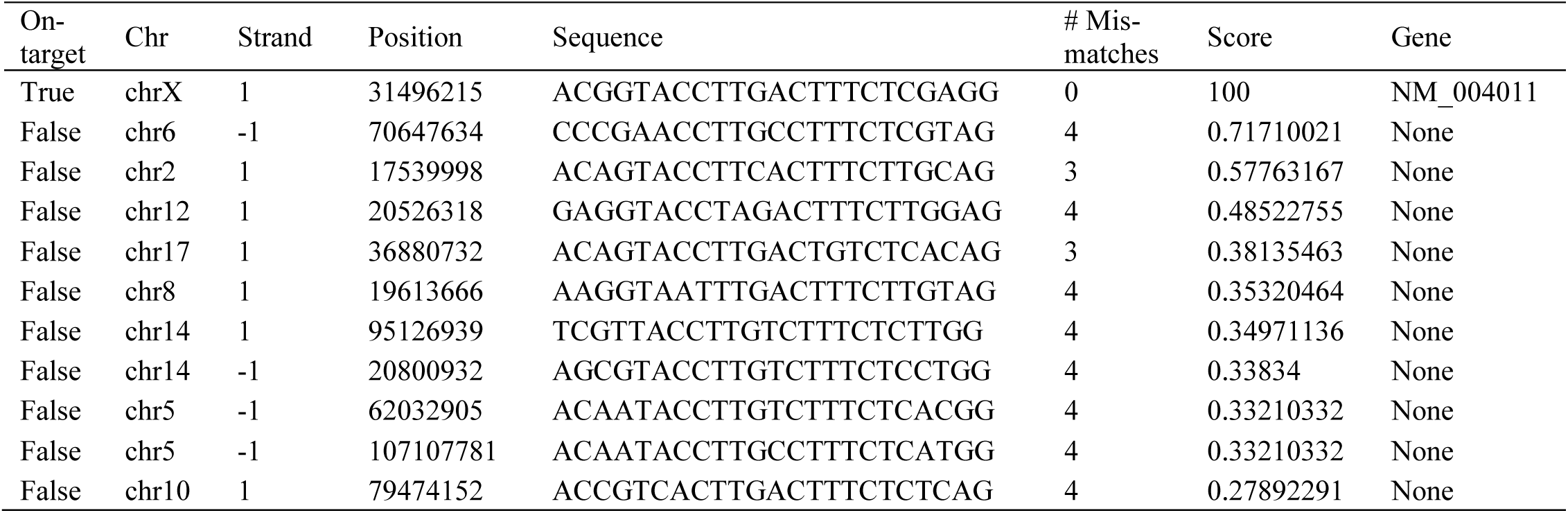
DMD-K2957fs sgRNA target sequence and predicted off-target sites.

**Table S3.** Gene Set Enrichment Analysis (DMD-K2957fs versus CORR-K2957fs, FDR < 0.05)

| **Day -1** |  |  |  |  |
| --- | --- | --- | --- | --- |
| **Gene set** | **pval** | **padj** | **ES** | **NES** |
| HALLMARK_EPITHELIAL_MESENCHYMAL_TRANSITION | 0.00234192 | 0.0234192 | 0.62363691 | 2.17331875 |
| HALLMARK_HYPOXIA | 0.00232019 | 0.0234192 | 0.5837144 | 1.99936245 |
| HALLMARK_COAGULATION | 0.00867679 | 0.04288165 | 0.57175214 | 1.75727054 |
| HALLMARK_COMPLEMENT | 0.00653595 | 0.03679176 | 0.50641651 | 1.69250891 |
| HALLMARK_UV_RESPONSE_DN | 0.00662252 | 0.03679176 | 0.50120265 | 1.68821864 |
| HALLMARK_GLYCOLYSIS | 0.00465116 | 0.03361345 | 0.46608141 | 1.60288321 |
| HALLMARK_ADIPOGENESIS | 0.00470588 | 0.03361345 | 0.46218475 | 1.59849062 |
| HALLMARK_WNT_BETA_CATENIN_SIGNALING | 0.00943396 | 0.04288165 | -0.6104547 | -1.6104716 |
| HALLMARK_G2M_CHECKPOINT | 0.0017331 | 0.0234192 | -0.6015427 | -2.0667642 |
| HALLMARK_MYOGENESIS | 0.0017331 | 0.0234192 | -0.614361 | -2.0763656 |
| HALLMARK_E2F_TARGETS | 0.00173611 | 0.0234192 | -0.6257252 | -2.1548396 |
| **Day 0** |  |  |  |  |
| **Gene set** | **pval** | **padj** | **ES** | **NES** |
| HALLMARK_EPITHELIAL_MESENCHYMAL_TRANSITION | 0.00381679 | 0.02385496 | 0.59092063 | 2.05977028 |
| HALLMARK_COMPLEMENT | 0.00358423 | 0.02385496 | 0.56808799 | 1.89806256 |
| HALLMARK_MITOTIC_SPINDLE | 0.00133869 | 0.01360544 | -0.490828 | -1.5450727 |
| HALLMARK_MYC_TARGETS_V1 | 0.00132979 | 0.01360544 | -0.5260148 | -1.664443 |
| HALLMARK_SPERMATOGENESIS | 0.00295858 | 0.02385496 | -0.6260033 | -1.7311382 |
| HALLMARK_G2M_CHECKPOINT | 0.00133511 | 0.01360544 | -0.6556942 | -2.0723203 |
| HALLMARK_E2F_TARGETS | 0.00132979 | 0.01360544 | -0.689547 | -2.1818998 |
| HALLMARK_MYOGENESIS | 0.00136054 | 0.01360544 | -0.724969 | -2.2554358 |
| **Day 2** |  |  |  |  |
| **Gene set** | **pval** | **padj** | **ES** | **NES** |
| HALLMARK_EPITHELIAL_MESENCHYMAL_TRANSITION | 0.00166113 | 0.00983671 | 0.71802554 | 2.63735759 |
| HALLMARK_COMPLEMENT | 0.00172414 | 0.00983671 | 0.60227869 | 2.10749818 |
| HALLMARK_UV_RESPONSE_DN | 0.0017094 | 0.00983671 | 0.58782135 | 2.07691829 |
| HALLMARK_INTERFERON_GAMMA_RESPONSE | 0.00172117 | 0.00983671 | 0.55923063 | 1.98385784 |
| HALLMARK_COAGULATION | 0.00180505 | 0.00983671 | 0.60939238 | 1.93814466 |
| HALLMARK_INTERFERON_ALPHA_RESPONSE | 0.00179211 | 0.00983671 | 0.59463371 | 1.91226973 |
| HALLMARK_TNFA_SIGNALING_VIA_NFKB | 0.00168067 | 0.00983671 | 0.50881113 | 1.82300241 |
| HALLMARK_APOPTOSIS | 0.0017331 | 0.00983671 | 0.514789 | 1.79951128 |
| HALLMARK_INFLAMMATORY_RESPONSE | 0.00178253 | 0.00983671 | 0.51490169 | 1.71871112 |
| HALLMARK_ALLOGRAFT_REJECTION | 0.00358423 | 0.01280082 | 0.48658315 | 1.60426556 |
| HALLMARK_APICAL_JUNCTION | 0.00508475 | 0.01412429 | 0.44816122 | 1.59601597 |
| HALLMARK_HYPOXIA | 0.005 | 0.01412429 | 0.42801344 | 1.54720303 |
| HALLMARK_CHOLESTEROL_HOMEOSTASIS | 0.01824818 | 0.03801703 | 0.48154909 | 1.515147 |
| HALLMARK_ESTROGEN_RESPONSE_LATE | 0.01034483 | 0.02586207 | 0.42302435 | 1.48228917 |
| HALLMARK_ESTROGEN_RESPONSE_EARLY | 0.01204819 | 0.02679118 | 0.41565868 | 1.47453965 |
| HALLMARK_XENOBIOTIC_METABOLISM | 0.01232394 | 0.02679118 | 0.42276588 | 1.46296512 |
| HALLMARK_P53_PATHWAY | 0.01145663 | 0.02679118 | 0.39584326 | 1.44249449 |
| HALLMARK_MITOTIC_SPINDLE | 0.00997506 | 0.02586207 | -0.3707878 | -1.4313851 |
| HALLMARK_WNT_BETA_CATENIN_SIGNALING | 0.0212766 | 0.04255319 | -0.5313992 | -1.5629091 |
| HALLMARK_MYC_TARGETS_V1 | 0.0049505 | 0.01412429 | -0.422513 | -1.6487597 |
| HALLMARK_OXIDATIVE_PHOSPHORYLATION | 0.00497512 | 0.01412429 | -0.4327252 | -1.6848361 |
| HALLMARK_SPERMATOGENESIS | 0.00217391 | 0.00983671 | -0.5476727 | -1.7802291 |
| HALLMARK_G2M_CHECKPOINT | 0.00247525 | 0.00983671 | -0.543074 | -2.11727 |
| HALLMARK_E2F_TARGETS | 0.00247525 | 0.00983671 | -0.5888373 | -2.297802 |
| HALLMARK_MYOGENESIS | 0.00255754 | 0.00983671 | -0.609027 | -2.3131645 |

Day 5
| Gene set | pval | padj | ES | NES |
| --- | --- | --- | --- | --- |
| HALLMARK_TNFA_SIGNALING_VIA_NFKB | 0.00119332 | 0.00335345 | 0.83605616 | 2.76632467 |
| HALLMARK_APOPTOSIS | 0.00121212 | 0.00335345 | 0.69566526 | 2.26954641 |
| HALLMARK_EPITHELIAL_MESENCHYMAL_TRANSITION | 0.0011655 | 0.00335345 | 0.66515294 | 2.23152291 |
| HALLMARK_P53_PATHWAY | 0.00116822 | 0.00335345 | 0.65412337 | 2.18570111 |
| HALLMARK_COMPLEMENT | 0.00121803 | 0.00335345 | 0.67067218 | 2.18197814 |
| HALLMARK_INFLAMMATORY_RESPONSE | 0.001287 | 0.00335345 | 0.69035559 | 2.16343309 |
| HALLMARK_HYPOXIA | 0.00117786 | 0.00335345 | 0.64300589 | 2.14206778 |
| HALLMARK_INTERFERON_GAMMA_RESPONSE | 0.00120482 | 0.00335345 | 0.64314571 | 2.11087735 |
| HALLMARK_KRAS_SIGNALING_UP | 0.00123609 | 0.00335345 | 0.63404723 | 2.04112252 |
| HALLMARK_REACTIVE_OXYGEN_SPECIES_PATHWAY | 0.00140845 | 0.00335345 | 0.72006118 | 2.04016119 |
| HALLMARK_IL2_STAT5_SIGNALING | 0.00121507 | 0.00335345 | 0.61922239 | 2.02919502 |
| HALLMARK_COAGULATION | 0.00134048 | 0.00335345 | 0.66716317 | 2.01904137 |
| HALLMARK_ESTROGEN_RESPONSE_EARLY | 0.00120482 | 0.00335345 | 0.61328155 | 2.0128598 |
| HALLMARK_ALLOGRAFT_REJECTION | 0.00130548 | 0.00335345 | 0.63547579 | 1.97190509 |
| HALLMARK_UV_RESPONSE_UP | 0.00122549 | 0.00335345 | 0.60887094 | 1.97090899 |
| HALLMARK_UV_RESPONSE_DN | 0.00120773 | 0.00335345 | 0.59882411 | 1.96010334 |
| HALLMARK_XENOBIOTIC_METABOLISM | 0.00123457 | 0.00335345 | 0.5811337 | 1.87506391 |
| HALLMARK_IL6_JAK_STAT3_SIGNALING | 0.0013947 | 0.00335345 | 0.63617743 | 1.83631147 |
| HALLMARK_ANDROGEN_RESPONSE | 0.00131579 | 0.00335345 | 0.58704122 | 1.81060743 |
| HALLMARK_ESTROGEN_RESPONSE_LATE | 0.00120919 | 0.00335345 | 0.52952307 | 1.72842335 |
| HALLMARK_MTORC1_SIGNALING | 0.00115207 | 0.00335345 | 0.50039332 | 1.6842422 |
| HALLMARK_UNFOLDED_PROTEIN_RESPONSE | 0.00376884 | 0.00856556 | 0.52084316 | 1.65486448 |
| HALLMARK_TGF_BETA_SIGNALING | 0.00977654 | 0.0181047 | 0.5699198 | 1.65477098 |
| HALLMARK_PROTEIN_SECRETION | 0.00522876 | 0.01136687 | 0.52660074 | 1.63450025 |
| HALLMARK_INTERFERON_ALPHA_RESPONSE | 0.01196809 | 0.02137158 | 0.51272088 | 1.5679143 |
| HALLMARK_CHOLESTEROL_HOMEOSTASIS | 0.01496599 | 0.02494331 | 0.51854236 | 1.54841017 |
| HALLMARK_ADIPOGENESIS | 0.01280559 | 0.0220786 | 0.42248509 | 1.41549634 |
| HALLMARK_MYOGENESIS | 0.02054795 | 0.03314185 | -0.3988695 | -1.5521062 |
| HALLMARK_MITOTIC_SPINDLE | 0.00740741 | 0.01526252 | -0.5100614 | -2.0177746 |
| HALLMARK_G2M_CHECKPOINT | 0.00775194 | 0.01526252 | -0.5715872 | -2.257701 |
| HALLMARK_E2F_TARGETS | 0.00793651 | 0.01526252 | -0.6228799 | -2.4496723 |

Day 7
| Gene set | pval | padj | ES | NES |
| --- | --- | --- | --- | --- |
| HALLMARK_INTERFERON_GAMMA_RESPONSE | 0.00111483 | 0.00437139 | 0.79491217 | 2.54616283 |
| HALLMARK_INTERFERON_ALPHA_RESPONSE | 0.00119617 | 0.00437139 | 0.8290613 | 2.48719542 |
| HALLMARK_EPITHELIAL_MESENCHYMAL_TRANSITION | 0.00108342 | 0.00437139 | 0.70054084 | 2.28095893 |
| HALLMARK_TNFA_SIGNALING_VIA_NFKB | 0.00110865 | 0.00437139 | 0.69843965 | 2.24927068 |
| HALLMARK_COAGULATION | 0.00122399 | 0.00437139 | 0.73878137 | 2.18716331 |
| HALLMARK_COMPLEMENT | 0.00113507 | 0.00437139 | 0.68255989 | 2.15401481 |
| HALLMARK_ALLOGRAFT_REJECTION | 0.00118765 | 0.00437139 | 0.67197649 | 2.04139381 |
| HALLMARK_INFLAMMATORY_RESPONSE | 0.00116822 | 0.00437139 | 0.66385028 | 2.03923343 |
| HALLMARK_APOPTOSIS | 0.00112486 | 0.00437139 | 0.64195017 | 2.0341882 |
| HALLMARK_IL6_JAK_STAT3_SIGNALING | 0.00131234 | 0.00437445 | 0.71882166 | 1.99293892 |
| HALLMARK_KRAS_SIGNALING_UP | 0.00114548 | 0.00437139 | 0.62573316 | 1.95530011 |
| HALLMARK_IL2_STAT5_SIGNALING | 0.00111483 | 0.00437139 | 0.57193951 | 1.82752538 |
| HALLMARK_P53_PATHWAY | 0.00109529 | 0.00437139 | 0.55672058 | 1.80323073 |
| HALLMARK_ESTROGEN_RESPONSE_EARLY | 0.00111483 | 0.00437139 | 0.54353914 | 1.74099629 |
| HALLMARK_UV_RESPONSE_DN | 0.00111982 | 0.00437139 | 0.53153163 | 1.69365913 |
| HALLMARK_CHOLESTEROL_HOMEOSTASIS | 0.00252845 | 0.00790139 | 0.57643751 | 1.67658878 |
| HALLMARK_ESTROGEN_RESPONSE_LATE | 0.00337838 | 0.00889047 | 0.50021491 | 1.58455321 |
| HALLMARK_APICAL_JUNCTION | 0.00330033 | 0.00889047 | 0.49013608 | 1.5768233 |
| HALLMARK_PROTEIN_SECRETION | 0.00714286 | 0.0170068 | 0.51662252 | 1.56632506 |
| HALLMARK_REACTIVE_OXYGEN_SPECIES_PATHWAY | 0.01874163 | 0.0334672 | 0.57441898 | 1.55963714 |
| HALLMARK_XENOBIOTIC_METABOLISM | 0.00685714 | 0.0170068 | 0.49227043 | 1.53911833 |
| HALLMARK_HYPOXIA | 0.00330396 | 0.00889047 | 0.47454221 | 1.53159678 |
| HALLMARK_ANDROGEN_RESPONSE | 0.01688782 | 0.03127373 | 0.50850786 | 1.52372299 |
| HALLMARK_UNFOLDED_PROTEIN_RESPONSE | 0.02186421 | 0.03644035 | 0.47794023 | 1.48507437 |
| HALLMARK_MYC_TARGETS_V1 | 0.01408451 | 0.02787068 | -0.3187556 | -1.3457584 |
| HALLMARK_G2M_CHECKPOINT | 0.01428571 | 0.02787068 | -0.3533847 | -1.4832261 |
| HALLMARK_MYC_TARGETS_V2 | 0.02531646 | 0.04083299 | -0.4166995 | -1.5123979 |
| HALLMARK_WNT_BETA_CATENIN_SIGNALING | 0.02090592 | 0.0360447 | -0.5159975 | -1.6556526 |
| HALLMARK_OXIDATIVE_PHOSPHORYLATION | 0.01449275 | 0.02787068 | -0.3957955 | -1.6661266 |
| HALLMARK_E2F_TARGETS | 0.01408451 | 0.02787068 | -0.4226466 | -1.7843773 |
| HALLMARK_MYOGENESIS | 0.01123596 | 0.02553626 | -0.4796555 | -2.0429474 |

**Table S4.**
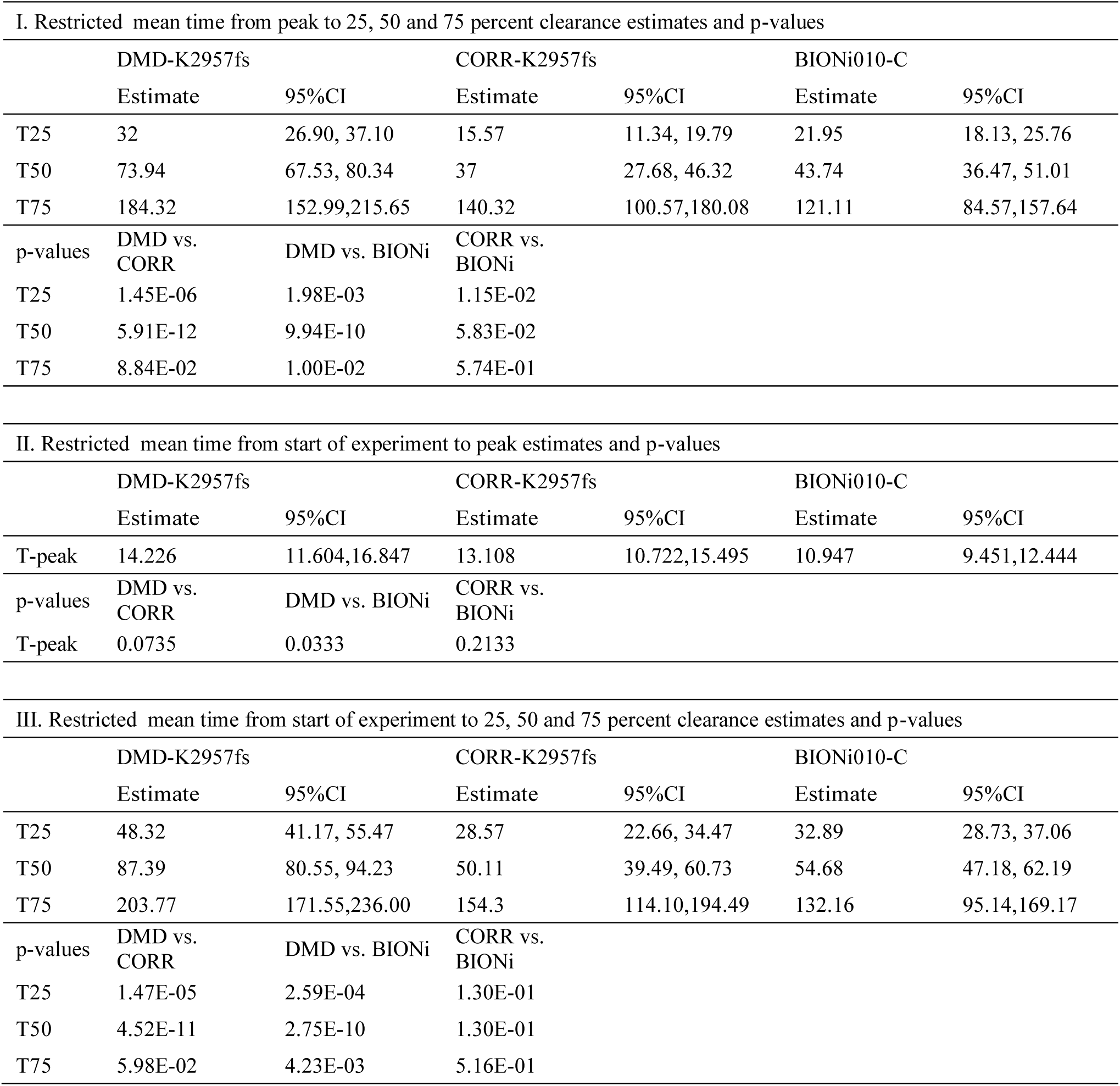
Restricted mean time estimates (sec) and p-values

**Table S5.** Exponential decay model parameters: 0-180 seconds and 0-360 seconds

| 0-180 sec | DMD-K2957fs |  | CORR-K2957fs |  | BIONi010-C |  |
| --- | --- | --- | --- | --- | --- | --- |
|  | Estimate | SE | Estimate | SE | Estimate | SE |
| Yo | 9.355 | 0.122 | 9.998 | 0.116 | 9.97 | 0.115 |
| Yf | 2.166 | 0.059 | 1.922 | 0.048 | 1.875 | 0.047 |
| Log(alpha) | -3.721 | 0.037 | -3.61 | 0.029 | -3.609 | 0.028 |

| 0-360 sec | DMD-K2957fs |  | CORR-K2957fs |  | BIONi010-C |  |
| --- | --- | --- | --- | --- | --- | --- |
|  | Estimate | SE | Estimate | SE | Estimate | SE |
| Yo | 9.193 | 0.094 | 9.832 | 0.092 | 9.809 | 0.091 |
| Yf | 1.983 | 0.022 | 1.75 | 0.02 | 1.709 | 0.019 |
| Log(alpha) | -3.815 | 0.022 | -3.692 | 0.018 | -3.688 | 0.018 |

(Units for Yo and Yf are concentration and unit for alpha is sec<sup>-1</sup>)

**Table S6.**
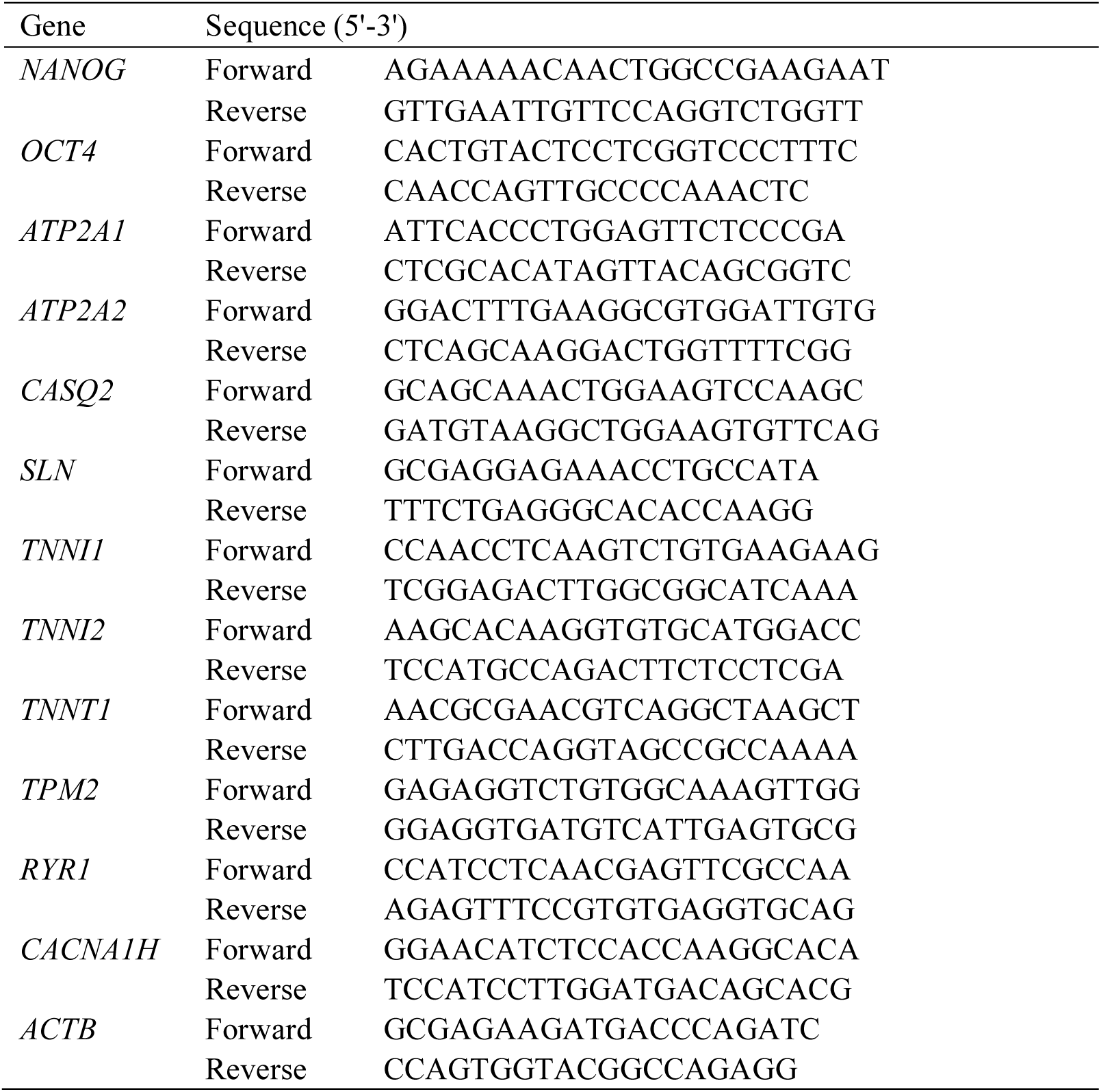
Primers for real-time quantitative PCR

Data S1. tpm_table.csv

Data S2. results_DMD_v_CORR_day_all time points.xlsx

Data S3. results_table_gsea_h.all.v6.2.entrez_all time points.xlsx

